# S1PR1 regulates the quiescence of lymphatic vessels by inhibiting laminar shear stress-dependent VEGF-C signaling

**DOI:** 10.1101/2020.02.27.968594

**Authors:** Xin Geng, Keisuke Yanagida, Racheal G. Akwii, Dongwon Choi, Lijuan Chen, YenChun Ho, Boksik Cha, Md. Riaj Mahamud, Karen Berman de Ruiz, Hirotake Ichise, Hong Chen, Joshua Wythe, Constantinos M. Mikelis, Timothy Hla, R. Sathish Srinivasan

## Abstract

During the growth of lymphatic vessels (lymphangiogenesis), lymphatic endothelial cells (LECs) at the growing front sprout by forming filopodia. Those tip cells are not exposed to circulating lymph, as they are not lumenized. In contrast, LECs that trail the growing front are exposed to shear stress, become quiescent and remodel into stable vessels. The mechanisms that coordinate the opposed activities of lymphatic sprouting and maturation remain poorly understood. Here we show that the canonical tip cell marker Delta-Like 4 (DLL4) promotes sprouting lymphangiogenesis by enhancing Vascular Endothelial Growth Factor C (VEGF-C) /VEGF Receptor 3 (VEGFR3) signaling. However, in lumenized lymphatic vessels laminar shear stress (LSS) inhibits the expression of DLL4, as well as additional tip cell markers. Paradoxically, LSS also upregulates VEGF-C/VEGFR3 signaling in LECs, but sphingosine 1-phosphate (S1P) receptor 1 (S1PR1) activity antagonizes LSS-mediated VEGF-C signaling to promote lymphatic vascular quiescence. Correspondingly, *S1pr1* loss in LECs induced lymphatic vascular hypersprouting and hyperbranching, which could be rescued by reducing *Vegfr3* gene dosage *in vivo*. In addition, S1PR1 regulates lymphatic vessel maturation by promoting membrane localization of the tight junction molecule Claudin-5. Our findings suggest a new paradigm in which LSS induces quiescence and promotes the survival of LECs by downregulating DLL4 and enhancing VEGF-C signaling, respectively. S1PR1 dampens LSS/VEGF-C signaling, thereby preventing sprouting from quiescent lymphatic vessels. These results also highlight the distinct roles that S1PR1 and DLL4 play in LECs when compared to their known roles in the blood vasculature.

## Introduction

Several signaling pathways (VEGF-A, Notch, BMP, PDGF-B, Angiopoietin-1 and −2, S1P, shear stress) are known to regulate angiogenesis in a highly coordinated manner (1). The migratory tip cells and the more-mature stalk cells have distinct characteristics (2). One of the important distinctions is that the tip cells are devoid of blood flow, whereas stalk cells actively experience hemodynamic cues and shear stress. DLL4, ADM, ESM1 and ANGPT2 are established markers of sprouting, angiogenic tip cells (3). The expression of at least some of these molecules is regulated by VEGF-A, which is expressed maximally in the vicinity of tip cells (4-7). However, to the best of our knowledge the mechanisms that restrict the expression of these molecules to a few cell layers close to the migrating front are not fully understood.

Growth of lymphatic vessels from pre-existing vessels is achieved via lymphangiogenesis, a process that is phenotypically similar to angiogenesis. The mechanisms that control lymphangiogenesis remain incompletely understood (8). VEGF-C binding to, and activation of, its cognate receptor VEGFR3 is the most well-studied pro-lymphangiogenic pathway known, and is necessary for the formation, migration and proliferation of lymphatic endothelial cells (LECs). Indeed, *Vegfc*^*-/-*^ mice lack LECs and *Vegfc*^*+/-*^ mice have severe lymphatic vessel hypoplasia (9). Likewise, mice harboring a dominant negative mutation in VEGFR3 feature hypoplastic lymphatic vessels (10). In contrast, VEGF-C overexpression in mice results in lymphatic vessel overgrowth and dysplasia (11). Consequently, mice overexpressing VEGF-C during a critical developmental time window develop lymphatic vascular defects, such as chylous ascites and chylothorax, and die rapidly after birth. Hence, a delicate balance of VEGF-C/VEGFR3 signaling is necessary for the proper patterning of the lymphatic vasculature. Specifically, while sprouts must form at the growing front under the influence of VEGF-C, the distal trailing vessels must remain stable and quiescent. Accordingly, negative regulators of VEGF-C/VEGFR3 signaling likely play key roles in coordinating productive lymphatic vascularization.

Shear stress produced by fluid flow is an important regulator of vascular development and physiology (12). Both oscillatory shear stress (OSS) and laminar shear stress (LSS) are necessary for lymphatic vascular development. OSS regulates the expression of molecules such as FOXC2 and GATA2 that are necessary for the maturation of lymphatic vessels and lymphatic valve development (13-16). In contrast, LSS regulates valve maturation (17) and promotes LEC proliferation by inhibiting Notch signaling (18, 19). However, whether any direct crosstalk exists between shear stress and VEGF-C signaling pathways is currently not known.

Sphingosine 1-phosphate (S1P) receptor 1 (S1PR1) is a G protein-coupled receptor (GPCR) that is necessary for preventing hypersprouting of the blood vessel endothelium. VEGF-A promotes the internalization and degradation of VE-cadherin from tight junctions to promote endothelial sprouting. S1PR1 antagonizes this process by stabilizing VE-cadherin assembly into adherens junctions and by inhibiting VEGF signaling thus limiting sprouting and enabling blood vessel maturation (20-22). Furthermore, S1PR1 is necessary for normal blood endothelial cell responses to LSS (23). Normally, in response to LSS blood endothelial cells align in the direction of blood flow and become quiescent (24). However, in the absence of S1PR1 blood endothelial cells fail to align in the direction of flow and do not activate LSS-responsive signals such as phosphorylation of downstream effector molecules like ERK, AKT and eNOS (23). Based on this knowledge, we hypothesized that S1PR1 also controls lymphatic vascular sprouting during lymphangiogenesis and maturation in response to LSS.

## RESULTS

### S1PR1 signaling is active in mature lymphatic vessels

Using our previously reported RNA-Sequencing (RNA-Seq) data, we determined that *S1PR1* is the most strongly expressed S1P receptor in primary cultured human LECs (HLECs) (**Supplementary Figure 1A**) (14, 25). Immunohistochemistry (IHC) on cryosections from embryonic day (E)17.5 wild-type mouse embryos confirmed that S1PR1 is expressed in LECs *in vivo* (**Supplementary Figure 1B**).

S1P binding to S1PR1 stimulates several signaling events such as G_i_-dependent Rac GTPase action and ß-arrestin recruitment to the plasma membrane, which leads to receptor internalization. To identify cells with S1PR1 signaling, we employed S1PR1-GFP reporter mice. These mice express: 1) S1PR1 C-terminally fused to a tetracycline transactivator (tTA), separated by a TEV protease recognition site, as well as 2) a ß-arrestin-TEV protease fusion protein. Thus, in the presence of S1P ligand, ß-arrestin-TEV protease is recruited to and cleaves the S1PR1-tTA chimeric receptor; free tTA then translocates to the nucleus to activate expression of a tetracyline response element-driven H2B-EGFP reporter (26). We analyzed cryosections of E16.5 S1PR1-GFP embryos and observed few GFP^+^ blood endothelial cells (**Supplementary Figure 1C**, arrowheads). Interestingly, most PROX1^+^ LECs were GFP^+^ (**Supplementary Figure 1C**, arrows), but some PROX1^+^ LECs were clearly GFP^-^ (**Supplementary Figure 1C**, red arrowheads).

To determine whether the GFP^+^ LECs were spatially restricted, we performed whole-mount IHC on the dorsal skin of E16.5 S1PR1-GFP embryos. Lymphatic vessels sprouting from the left and right lymph sacs migrate towards the midline of the dorsal skin in a stereotypic manner (27). These lymphatic vessels featured elevated GFP reporter expression compared to blood vessels (**Figure 1A**, compare white and yellow arrowheads, respectively). Furthermore, GFP was evident in maturing lymphatic vessels (**Figure 1A**, white arrowheads) and in lymphatic valves (**Figure 1A**, green arrowheads), but LECs at the migrating front lacked GFP expression (**Figure 1A**, arrows). As expected, GFP was not expressed in single transgenic H2B-GFP negative control littermates (**Figure 1A**). Thus S1PR1 signaling is restricted to quiescent LECs in lymphatic vessel stalks, rather than tip LECs undergoing active lymphangiogenesis.

**Figure 1:**
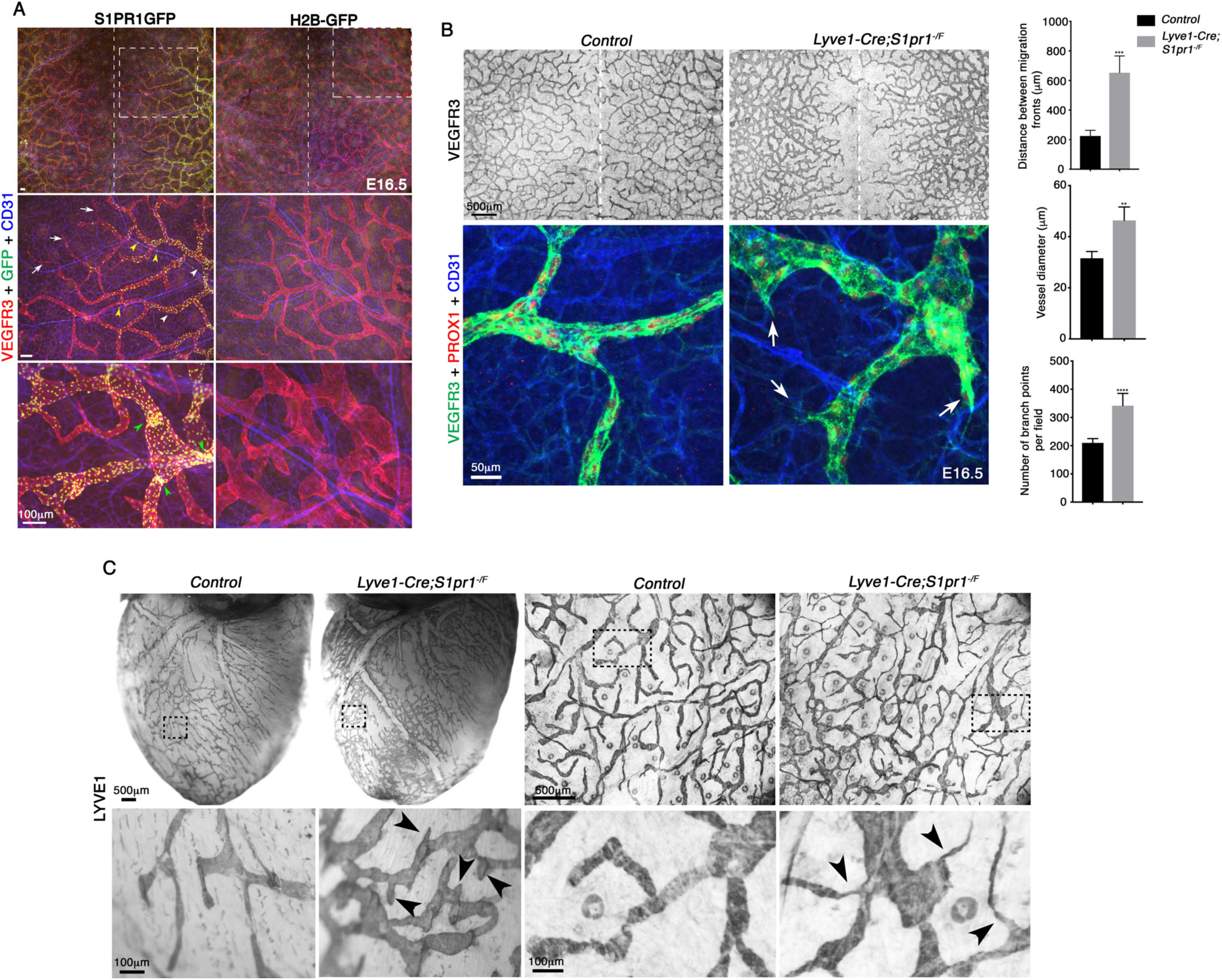
S1PR1 inhibits lymphatic vascular hypersprouting. *(A) S1PR1 signaling is active in stable lymphatic vessels and in lymphatic valves*. In the dorsal skin VEGFR3^+^ lymphatic vessels migrate from the two sides towards the midline (dotted line). H2B-GFP embryos were devoid of GFP expression. Few GFP^+^ cells were observed in CD31^+^ blood vessels (yellow arrowheads) or in the migrating front of lymphatic vessels (arrows) in S1PR1-GFP mice. In contrast, LECs that were behind the migrating front (arrowheads) and lymphatic valves (green arrowheads) were predominantly GFP^+^. Statistics: n=4 for both S1PR1GFP and H2B-GFP. (*B, C*) *Deletion of* S1pr1 *from LECs results in lymphatic vascular hypersprouting*. (B) The lymphatic vessels of E16.5 embryos lacking *S1pr1* were dilated, had excessive number of branches and did not migrate completely to the midline. Additionally, while the control embryos were devoid of sprouts in between the lymphatic vessel branches, several sprouts were observed in embryos lacking *S1pr1* (arrows). Statistics: n=3 for control embryos; n=6 for mutant embryos. (C) Heart and ear skin of adult mice lacking S1PR1 had higher lymphatic vessel density and excessive number of sprouts (arrowheads). Statistics: n=6 for control animals; n=4 for mutant animals.

### S1PR1 prevents lymphatic vessel hypersprouting and hyperbranching

To investigate the function of S1PR1 signaling in the lymphatic endothelium, we used *Lyve1-Cre* (28) to delete *S1pr1* (29) specifically in LECs of the dorsal skin vasculature. We generated *Lyve1-Cre;S1pr1*^*f/-*^ embryos by breeding *Lyve1-Cre;S1pr1*^*+/-*^ with *S1pr1*^*f/f*^ mice. E16.5 *Lyve1-Cre;S1pr1*^*f/-*^ embryos did not display any obvious defects such as edema or blood-filled lymphatic vasculature suggesting that the lymphatic drainage is happening in mutants (data not shown). However, analysis of the dorsal skin by whole mount immunohistochemistry revealed that the E16.5 *Lyve1-Cre;S1pr1*^*f/-*^ embryos displayed a higher number of lymphatic vessel branches (**Figure 1B**). The *S1pr1* mutant lymphatic vessels were dilated and did not reach the dorsal midline, unlike control vessels (**Figure 1B**). The migrating front appeared indistinguishable between control and mutant embryos (data not shown). In contrast, excessive sprouts were seen within the lymphatic plexus of mutant embryos (**Figure 1B**, arrows).

We genotyped over 400 animals at postnatal day 10 (P10), but obtained only 4 *Lyve1-Cre;S1pr1*^*f/-*^ mice, indicating perinatal lethality. The surviving adult animals displayed increased lymphatic vascular sprouts in the heart and the ears (**Figure 1C**, arrowheads). These results suggest that S1PR1 prevents lymphatic vascular hypersprouting and hyperbranching during embryogenesis and organogenesis.

### S1PR1 regulates the expression of claudin-5 in maturing lymphatic vessels

During angiogenesis, S1PR1 regulates the stability of quiescent blood endothelial cells by promoting the VE-Cadherin assembly into adherens junctions (20, 23). VE-Cadherin is also a mechanosensory molecule that is critical for sensing shear stress (30). Mice lacking *Cdh5* (which encodes VE-Cadherin) in blood endothelial cells recapitulated the hypersprouting phenotype of mice lacking *S1pr1* (20). We did not observe any obvious reduction in VE-Cadherin expression within the lymphatic vessels of E17.5 *Lyve1-Cre;S1pr1*^*f/-*^ embryos, although VE-Cadherin appeared to be mislocalized in some LECs (**Figure 2A**, red arrowheads). In addition, lymphatic vascular hypersprouting and hyperbranching were not reported in *Cdh5* lymphatic mutant mice (31, 32). Hence, we conclude that VE-Cadherin expression is not regulated by S1PR1 in LECs although its localization appears to be regulated.

**Figure 2:**
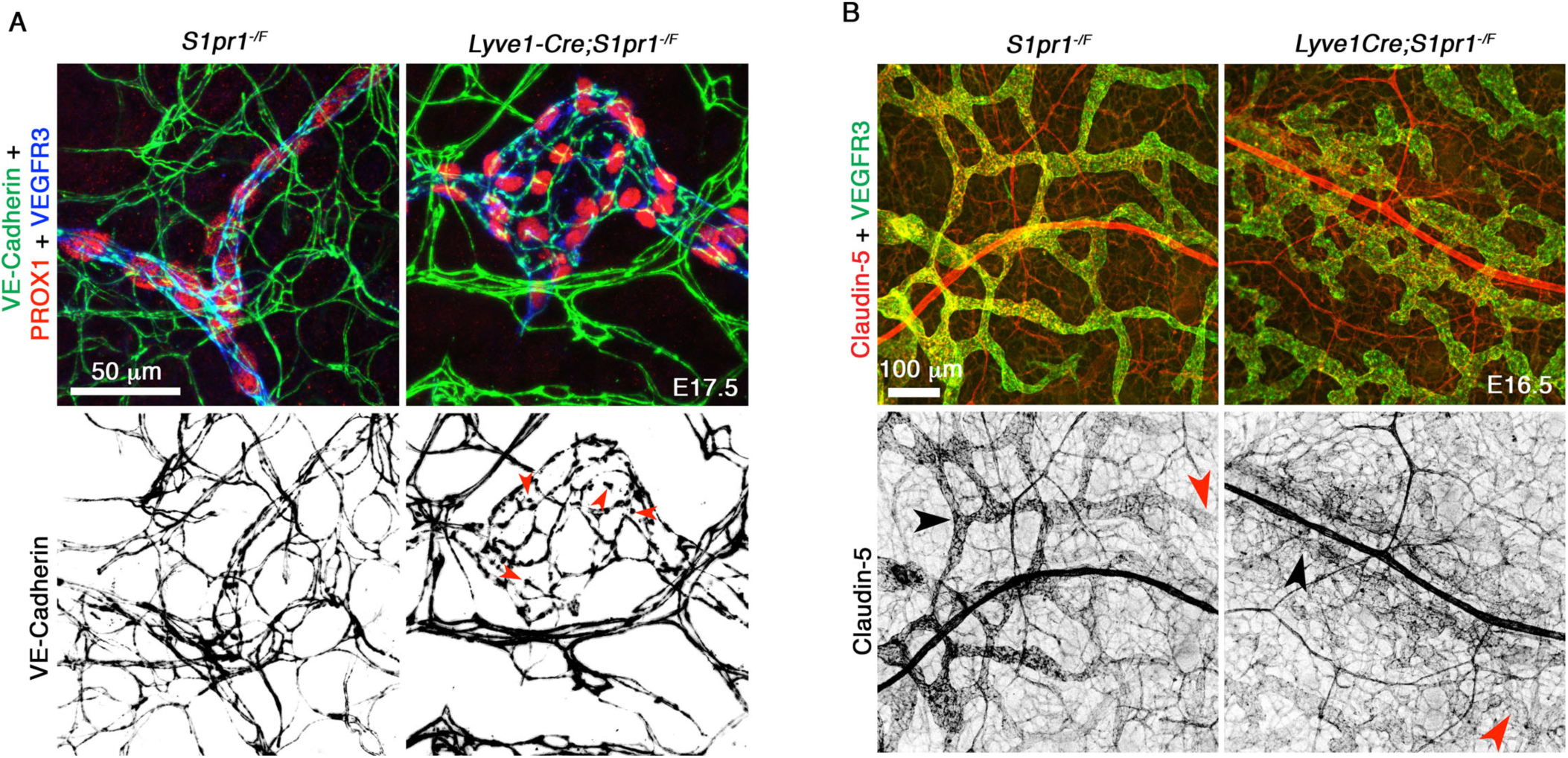
S1PR1 regulates claudin-5 expression in the developing lymphatic vessels. *(A) VE-Cadherin expression was modestly defective in the lymphatic vessels of embryos lacking S1PR1*. VE-Cadherin was uniformly expressed with “zipper-like” structure in the lymphatic vessels of E17.5 control embryos. Breaks in VE-Cadherin localization were observed (red arrowheads) at restricted locations within the lymphatic vessels of embryos lacking S1PR1. Statistics: n=6 for control embryos; n=3 for mutant embryos. *(B) Claudin-5 was severely downregulated in embryos lacking S1PR1*. In control E16.5 embryos claudin-5 was expressed in a gradient manner along the developing lymphatic vessels with weaker expression at the growing tips (red arrows) and stronger expression in stable lymphatic vessels behind the migrating front (black arrows). In contrast, expression of claudin-5 was uniformly low in embryos lacking S1PR1. Statistics: n=6 for control embryos; n=10 for mutant embryos.

S1PR1 maintains blood-brain barrier by regulating the proper localization of tight junction proteins including Claudin-5 (33). Therefore, we tested whether Claudin-5 localization could be defective in the lymphatic vessels of *Lyve1-Cre;S1pr1*^*f/-*^ embryos. Intriguingly, Claudin-5 was expressed in a gradient within the growing lymphatic vessels at E16.5. Claudin-5 expression was lower in the migrating lymphatic vessels compared to the patent vessels of the posterior plexus (**Figure 2B**, compare red and black arrowheads, respectively). This gradient of Claudin-5 expression was reminiscent of reporter activity in the S1PR1-GFP embryos (**Figure 1A**). Therefore, we investigated whether expression of Claudin-5 is regulated by S1PR1. Claudin-5 expression remained low in the migrating lymphatic front of *Lyve1-Cre;S1pr1*^*-/f*^ embryos (**Figure 2B**, red arrowhead). However, Claudin-5 was dramatically downregulated in the posterior lymphatic plexus (**Figure 2B**, black arrowhead). This observation indicated that S1PR1 regulates the expression of Claudin-5 in the developing lymphatic vasculature.

In summary, our data suggest that S1PR1 regulates the quiescence and maturation of lymphatic vasculature by preventing hypersprouting and by promoting cell-junction formation respectively.

### DLL4 is a pro-lymphangiogenic molecule in vitro and in vivo

S1PR1 activity is excluded from the migrating tip cells (**Figure 1A**) and deletion of *S1pr1* from LECs resulted in the dramatic increase in the number of tip cells (**Figure 1B**). Delta-like 4 (DLL4) is the prototypic tip-cell marker in blood vasculature. *Dll4* encodes a Notch ligand that is critical for blood vascular patterning. VEGF-A activation of VEGFR2 in tip cells increases *Dll4* expression (5). In turn, DLL4 activates Notch signaling in adjacent stalk cells, which decreases VEGFR2 levels and inhibits excessive sprouting (5, 34, 35). Deletion of just one allele of *Dll4* results in embryonic death due to arteriovenous malformations and hypersprouting of the blood endothelial cells (5, 34-37). DLL4 inhibits VEGF-C signaling *in vivo* in the zebrafish blood vasculature (38). As in blood vessels, DLL4 is expressed in the tip cells of intestinal lacteals (39). Likewise, DLL4 is expressed in the growing front of dermal lymphatic vessels (**Supplementary Figure 2**). Both pro- and anti-lymphangiogenic properties are attributed to DLL4. Overexpression of DLL4-Fc, which inhibits the activity of endogenous DLL4, results in ectopic sprouts (40). In contrast, *Dll4* is required for postnatal lymphangiogenesis during wound healing and intestinal lacteal regeneration (39, 41). Given that the angiogenic phenotype of *Dll4*^*+/-*^ mice is similar to loss of S1PR1, we investigated whether *Dll4* is required for normal lymphatic vessel patterning during mouse development.

IHC revealed that the lymphatic vessels in the dorsal skin of E16.5 *Dll4*^*+/-*^ embryos were severely hypoplastic (**Figure 3A**). Additionally, *Dll4*^*+/-*^ mutant embryo lymphatic vessels were less migratory than control littermates and had significantly fewer branch points (**Figure 3A**). These results indicate that in contrast to its role in blood vasculature, DLL4 promotes the growth of lymphatic vessels. VEGF-C and its receptor VEGFR3 are critical, established positive regulators of lymphangiogenesis. Hence, by using in vitro assays we tested whether DLL4 regulates VEGF-C signaling. Briefly, we transfected HLECs with *siDLL4*, then treated them with VEGF-C. As expected, HLECs treated with *siDLL4* showed downregulation of DLL4 and Notch1 activation, as assessed by presence of the cleaved Notch1 intracellular domain (NICD) (**Figure 3B**). Consistent with the pro-lymphangiogenic role of DLL4 in vivo, siDLL4 treatment reduced the levels of pERK and pAKT in HLECs treated with VEGF-C (**Figure 3B**). These results suggest that DLL4 enhances VEGF-C signaling in tip LECs to promote lymphangiogenesis.

**Figure 3:**
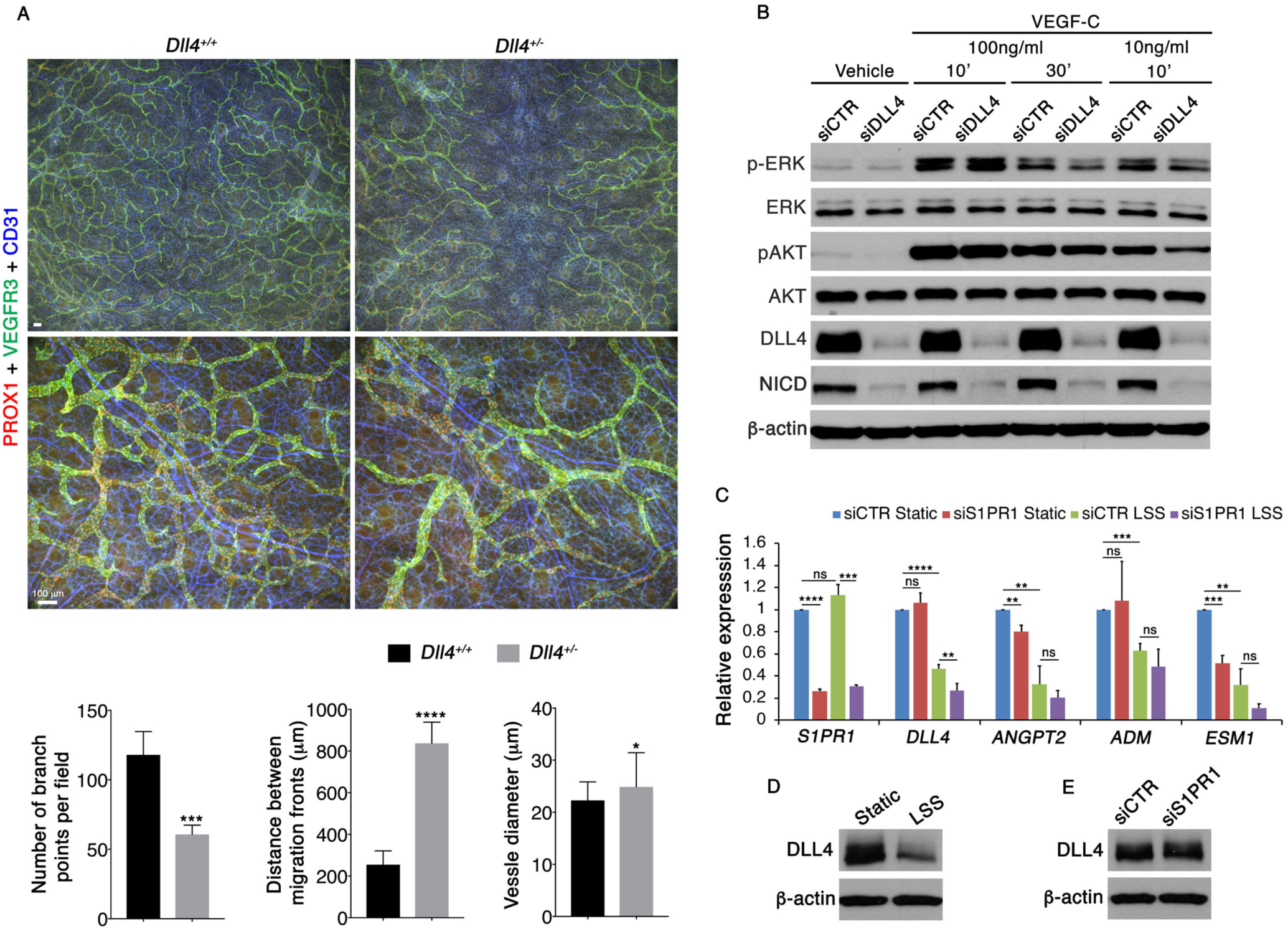
LSS inhibits the expression of pro-lymphangiogenic tip cell molecules in a S1PR1 independent manner. *(A) Lymphatic vessels of Dll4*^*+/-*^ *mice are hypoplastic*. Lymphatic vessels in the dorsal skin of E16.5 *Dll4*^*+/-*^ mice had fewer branch points, had migrated lesser distance and were mildly dilated when compared to their wild type littermates. *(B) DLL4 enhances VEGF-C signaling*. Signaling pathway activated by VEGF-C was investigated using HLECs that were transfected with either control siRNA or siDLL4. Phosphorylation of ERK and AKT was reduced in HLECs transfected with siDLL4. Statistics: n=4 for both wild type and *Dll4*^*+/-*^ embryos. *(C-E) DLL4, ANGPT2, ADM and ESM1 are inhibited by LSS*. HLECs transfected with control siRNA or siS1PR1 were cultured under LSS for 24 hours and the expression of tip cell markers was quantified by qRT-PCR (C) or by Western blot (D, E). (C) LSS inhibited the expression of all tip cell markers except *DLL4* in an S1PR1-independent manner. *DLL4* was slightly but significantly more reduced in siS1PR1 transfected HLECs. (D, E) While LSS reduced the expression of DLL4 (D), no obvious differences were was observed between control siRNA and siS1PR1 transfected HLECs that were cultured under LSS (E). Statistics: n=3 for all experiments. **p < 0.01, ***p < 0.001, *** p < 0.0001. Error bars in graphs represent ±SEM.

### Laminar Shear Stress (LSS) inhibits the expression of tip-cell expressed molecules in an S1PR1-independent manner

In addition to DLL4, Angiopoietin-2 (ANGPT2), Endothelial Cell Specific Marker 1 (ESM1), and Adrenomedullin (ADM) are expressed in blood endothelial tip cells (3). Both ANGPT2 and ADM promote lymphatic vascular growth and patterning *in vivo* (42, 43), while ESM1 can enhance HLEC proliferation induced by VEGF-C *in vitro* (7). Thus, all tip-cell enriched genes are necessary for lymphatic vascular growth. Furthermore, tip cells are not exposed to shear stress as they lack a functional lumen (2, 22).

We exposed HLECs to 5 dynes/cm^2^ LSS as described by Choi et al (18, 19) and investigated whether S1PR1 and LSS could synergize in restricting the expression of tip cell markers. Indeed, LSS potently inhibited the expression of *DLL4, ANGPT2, ESM1* and *ADM* in both control siRNA and *siS1PR1*-treated HLECs (**Figure 3C**). Knockdown of *S1PR1* slightly reduced the expression of *ANGPT2* in HLECs cultured under static conditions (**Figure 3C**). Expression of *ESM1* was also potently downregulated by *siS1PR1* (**Figure 3C**). However, no obvious differences in expressions of *ESM1, ANGPT2* or *ADM* were observed between control siRNA and *siS1PR1* transfected HLECs cultured under LSS (**Figure 3C**). A slight reduction in *DLL4* was observed in *siS1PR1* transfected HLECs cultured under LSS (**Figure 3C**). However, while Western blotting confirmed the downregulation of DLL4 expression by LSS (**Figure 3D**), we did not observe an obvious difference in DLL4 expression between control siRNA and *siS1PR1* treated HLECs that were cultured under LSS (**Figure 3E**). These results indicate that LSS inhibits the expression of several tip-cell molecules in HLECs in an S1PR1-independent manner.

### S1PR1 does not regulate VEGF-C signaling in HLECs cultured under static conditions

Having excluded the role of S1PR1 in regulating tip cell identity we investigated whether S1PR1 signaling could directly regulate VEGF-C signaling or LSS in HLECs. Briefly, HLECs were serum starved for 6 hours, pretreated with either the S1PR1 agonist SEW2871, or the S1PR1 antagonist W146 for 1 hour, and then treated with 100 ng/ml VEGF-C. We observed no obvious differences in either phosphorylated ERK or AKT between control and SEW2871-treated HLECs, whereas W146 treatment slightly reduced the levels of pERK and pAKT (**Figure 4A**). In addition, HLECs treated with control *siRNA* or *siS1PR1* for 48 hours prior to stimulation with either 100 ng/ml or 10 ng/ml VEGF-C did not show any obvious differences in pERK or pAKT levels (**Figure 4B** and data not shown). Together, these data suggest that S1PR1 signaling does not inhibit VEGF-C signaling in HLECs under static conditions.

**Figure 4:**
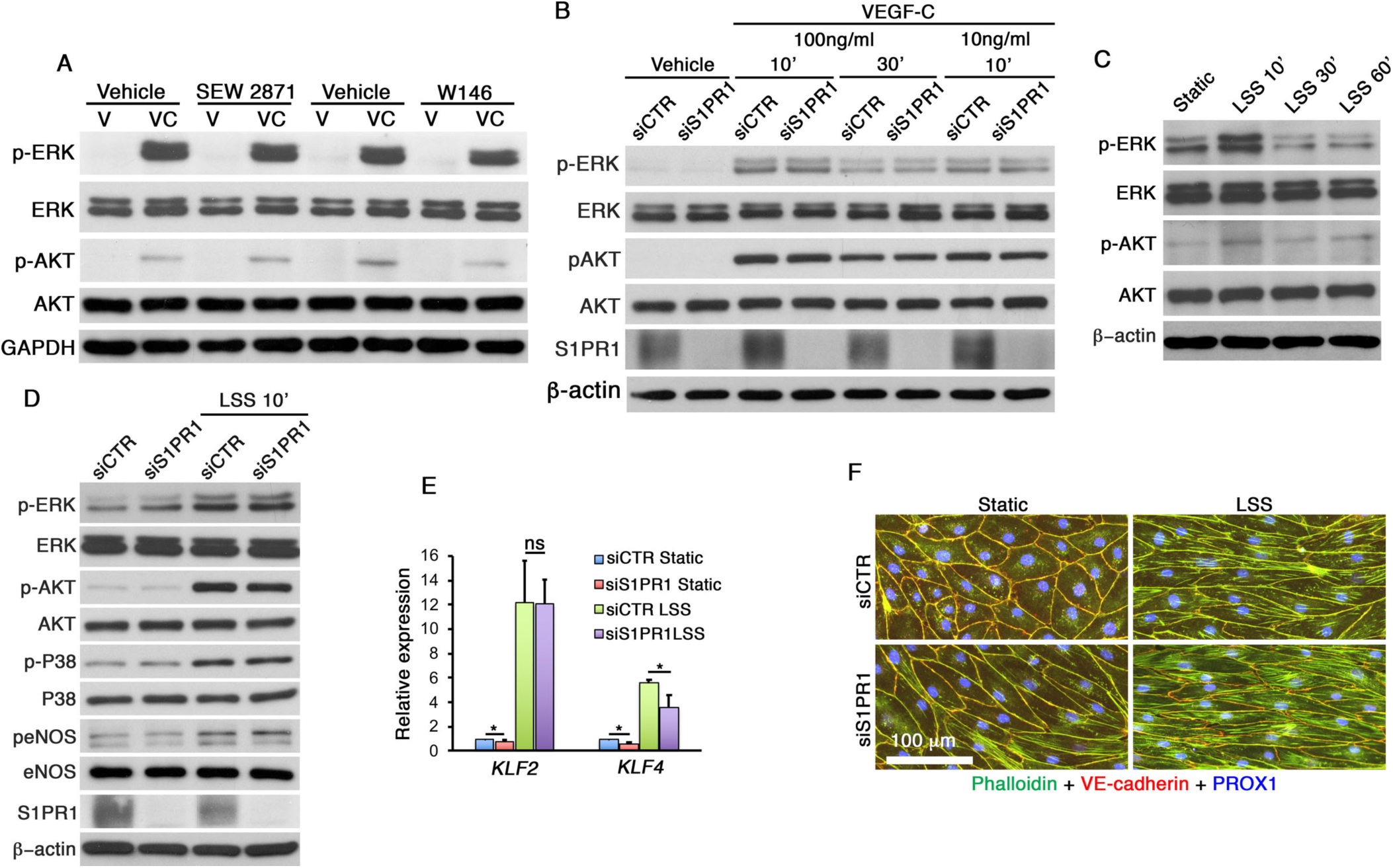
S1PR1 does not inhibit VEGF-C signaling in statically cultured HLECs, and it is not necessary for canonical LSS response. *(A, B) S1PR1 does not regulate VEGF-C signaling in statically cultured HLECs*. (A) HLECs were treated with 100 ng/ml VEGF-C in the presence or absence of S1PR1 agonist SEW 2871 or antagonist W146. SEW 2871 did not cause any obvious differences in VEGF-C signaling (phosphorylation of ERK and AKT). W146 slightly reduced the VEGF-C signaling. (B) HLECs were transfected with control siRNA or siS1P1 and then treated with the indicated concentrations of VEGF-C. No obvious differences were observed in VEGF-C signaling between control and siS1PR1 treated HLECs. Statistics: n=3 for A; n=4 for B. *(C-F) S1PR1 does not regulate canonical LSS-responses in HLECs*. (C) Exposure of HLECs to LSS (5 dynes/cm^2^) for 10 minutes activated the phosphorylation of ERK and AKT. Prolonged exposure (30-60 minutes) to LSS caused the phosphorylation of ERK and AKT to return to background levels (D) LSS induced phosphorylation of ERK, AKT, P38 and eNOS was not affected by siS1PR1. (E) Shear stress responsive genes *KLF2* and *KLF4* were mildly reduced in siS1PR1 treated HLECs cultured under static condition. Both *KLF2* and *KLF4* were upregulated in both control and siS1P1 treated HLECs that were cultured with LSS for 24 hours. A slight reduction in *KLF4* expression was observed in siS1PR1 treated HLECs compared to controls. (F) siS1PR1 treatment caused the elongation of statically cultured HLECs. However, both control and siS1PR1 transfected HLECs align normally in the direction of LSS. Statistics: n=3 for C, E and F; n=4 for D. *p < 0.05. Error bars in graphs represent ±SEM.

### S1PR1 is not necessary for the LSS response in HLECs

LSS promotes the quiescence of blood vascular endothelial cells (44, 45). S1PR1 is necessary for HUVEC alignment in response to LSS and for activating LSS-regulated signaling pathways (MEK/ERK, PI3K/AKT, and eNOS) (23). Likewise, LSS elevated pERK and pAKT levels in HLECs within 10 minutes, and this increase returned to baseline levels after 30 minutes (**Figure 4C**). To determine whether S1PR1 is required for LSS-mediated activation of flow-induced signaling pathways, we exposed HLECs transfected with control *siRNA* and *siS1PR1* to LSS. Phosphorylation of ERK, AKT, p38 and eNOS was indistinguishable between control *siRNA* and *siS1PR1*-treated HLECs (**Figure 4D**). Furthermore, both control and *siS1PR1*-treated HLECs exposed to 24 hours of LSS had upregulated the shear stress-responsive genes *KLF2* and *KLF4* (**Figure 4E**), and were elongated and aligned in the direction of fluid flow (**Figure 4F**). Altogether, these results demonstrate that that S1PR1 is not necessary for canonical LSS responses in HLECs.

### LSS enhances VEGF-C signaling, which is antagonized by S1PR1

LSS sensitizes blood endothelial cells to BMP9 and BMP10 signaling (46). To test whether LSS sensitizes HLECs to VEGF-C stimulation, we exposed HLECs to LSS for 24 hours, and then treated with 10 ng/ml, 5 ng/ml or 1 ng/ml of VEGF-C for 10 minutes. We found that LSS enhanced the phosphorylation of ERK and AKT at all tested concentrations of VEGF-C compared to static culture conditions (**Figure 5A**). Thus, LSS potentiates VEGF-C signaling in HLECs.

**Figure 5:**
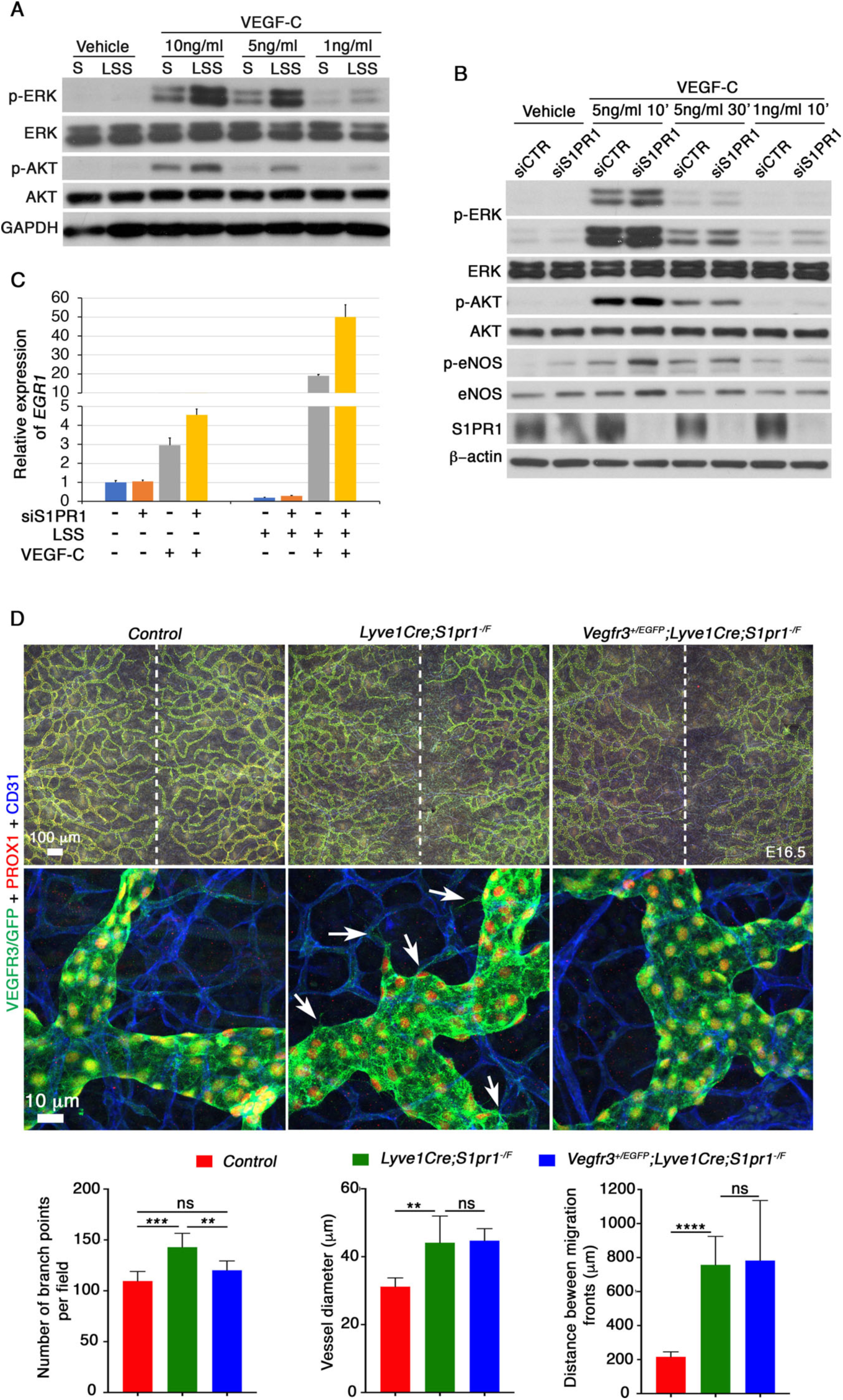
S1PR1 inhibits LSS-enhanced VEGF-C signaling. *(A) Exposure to LSS enhances VEGF-C signaling in HLECs*. HLECs were cultured with 5 dynes/cm^2^ of LSS for 24 hours following which they were treated with VEGF-C. Phosphorylation of ERK and AKT was enhanced in LSS-exposed HLECs compared to statically cultured cells (S). Statistics: n=4. *(B, C) S1PR1 antagonizes LSS enhanced VEGF-C signaling*. HLECs were transfected with control siRNA or siS1PR1, exposed to LSS for 24 hours and then treated with VEGF-C for (B) 10 or 30 minutes under static culture condition or for (C) 1 hour in the presence of LSS. (B) Phosphorylation of ERK and eNOS, but not AKT, was enhanced in siS1PR1 transfected HLECs. Total eNOS was also increased in siS1PR1 transfected HLECs. (C) VEGF-C induced expression of *EGR1* was dramatically enhanced by LSS and it was further enhanced by siS1PR1. Statistics: n=4 for B;C is representative of 3 independent experiments. Results from the other 2 experiments are presented in Supplementary Figure 3. P values were not presented due to large variability in fold activation of *EGR1* between independent experiments *(D) Heterozygosity of Vegfr3 rescues the hypersprouting phenotype in embryos lacking S1P1*. E16.5 *Lyve1-Cre; S1P1*^*-/f*^ embryos were generated in the *Vegfr3*-heterozygous background. Increased number of branch points that were observed in *Lyve1-Cre; S1P1*^*-/f*^ embryos was rescued by the loss of one allele of *Vegfr3*. Additionally, the excessive number of sprouts that were seen in *Lyve1-Cre; S1P1*^*-/f*^ embryos (arrows) was also reduced. However, the lymphatic vessels remained enlarged and they did not migrate completely to the midline. Statistics: n=5 for control embryos; n=7 for *Lyve1-Cre; S1P1*^*-/f*^ embryos; n=5 for *Vegfr3*^*+/EGFP*^;*Lyve1-Cre; S1P1*^*-/f*^ embryos. **p < 0.01, ***p < 0.001, *** p < 0.0001. Error bars in graphs represent ±SEM.

To determine whether S1PR1 affects this process, we transfected HLECs with control siRNA or siS1PR1, exposed them to LSS and then treated with VEGF-C. We found that phosphorylation of ERK and eNOS was increased in *siS1PR1*-treated HLECs compared to control cells (**Figure 5B**). Total eNOS was also increased in *siS1PR1*-treated HLECs. In contrast, AKT phosphorylation was unchanged. These data suggest that S1PR1 antagonizes specific downstream pathways of LSS-enhanced VEGF-C signaling in HLECs.

*Early growth response 1 (EGR1)* is an immediate early gene that is activated by VEGF-C in HLECs (4), so we examined whether this process is affected by LSS and S1PR1. *EGR1* expression was downregulated by LSS (**Figure 5C**). Although VEGF-C induced *EGR1* expression in both statically cultured and LSS-treated HLECs, the fold induction was higher in HLECs grown under LSS (**Figure 5C** and **Supplementary Figure 3**). Importantly, *EGR1* was upregulated even further in siS1PR1-treated HLECs compared to control cells that were cultured under LSS. Together these results further suggest that S1PR1 signaling antagonizes LSS-enhanced VEGF-C signaling.

### *Vegfr3*-heterozygosity ameliorates hypersprouting caused by the deletion of *S1pr1*

Given our findings that S1PR1 inhibits VEGF-C/VEGFR3 signaling *in vitro*, we hypothesized that genetic reduction of VEGF-C/VEGFR3 signaling components would ameliorate the phenotypes observed in *Lyve1-Cre; S1pr1*^*-/f*^ mice. Indeed, *Vegfr3* heterozygosity rescued the increased numbers of sprouts and branches that were observed in lymphatic *S1pr1* mutants (arrows) (**Figure 5D**). However, neither the increased vessel diameter, nor delayed migration was rescued by *Vegfr3*-heterozygosity. Furthermore, Claudin-5 expression remained reduced in *Lyve1-Cre;S1pr1*^*f/-*^;*Vegfr3*^*+/-*^ embryos (**Supplementary Figure 4**) suggesting that these *S1PR1* mutant phenotypes are likely not dependent on VEGF-C/VEGFR3 signaling.

In summary, LSS plays a dual role in regulating lymphangiogenesis. On the one hand LSS inhibits the expression pro-lymphangiogenic molecules such as DLL4, ANGPT2, ESM1 and ADM, and on the other hand LSS promotes VEGF-C signaling. S1PR1 is not involved in LSS-mediated downregulation of pro-lymphangiogenic molecules. However, S1PR1 antagonizes LSS-enhanced VEGF-C signaling.

### S1PR1 regulates the membrane localization of Claudin-5 by inhibiting RhoA

To investigate how S1PR1 regulates Claudin-5, we treated cultured HLECs with the S1PR1 inhibitor W146. 30 minutes of W146 treatment dramatically decreased Claudin-5 staining in the periphery of HLECs (**Figure 6A**). However, Western blotting revealed that the expression of claudin-5 was not downregulated in W146-treated HLECs (data not shown), suggesting that S1PR1 inhibition causes mislocalization of Claudin-5. No obvious changes were observed in the expression of other tight junction and adherens junction proteins ZO-1 and VE-Cadherin, although they were expressed with a discontinuous (zigzag) pattern in W146-treated HLECs (**Figure 6A** and **Supplementary Figure 5A**, respectively). Furthermore, gaps could be seen in between cells, suggesting that the cell-cell interactions were compromised (**Supplementary Figure 5A**, white arrowheads). We used *in vitro* cell permeability assays to verify this observation. Briefly, we coated plates with biotin-conjugated gelatin before seeding HLECs. Confluent HLECs were treated with W146 for 30 minutes, fixed, washed and stained with Alexa 488-conjugated streptavidin. Intercellular gaps were revealed by a green signal. As shown in **Supplementary Figure 6**, inhibition of S1PR1 by W146 resulted in increased intercellular gaps.

**Figure 6:**
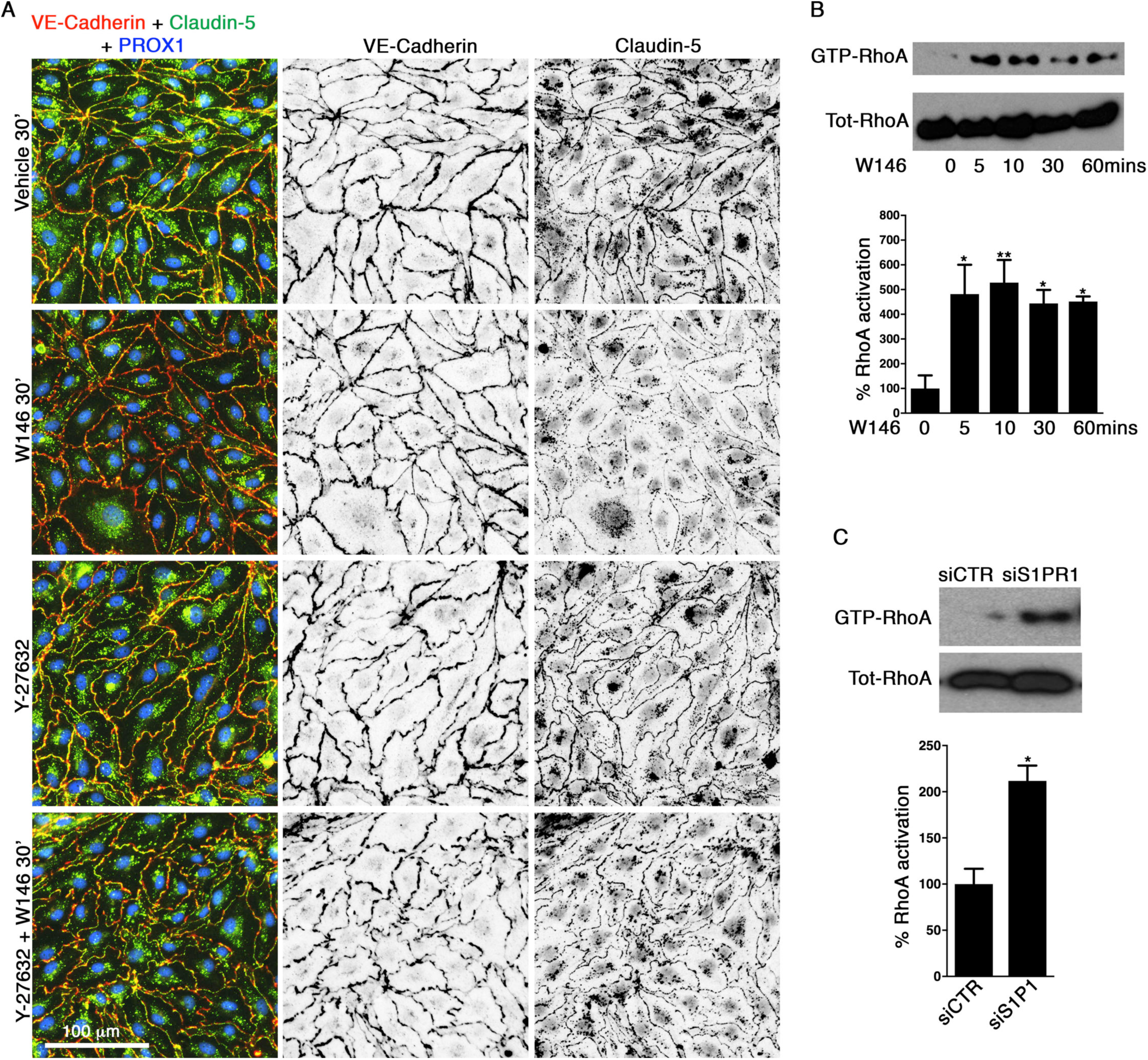
S1PR1 inhibits RhoA signaling to maintain Claudin-5 expression in HLECs. *(A) S1PR1 regulates the cell-membrane localization of claudin-5 by inhibiting RhoA/ROCK signaling*. VE-Cadherin and claudin-5 were localized to the cell membrane in vehicle-treated HLECs. W146 treatment for 30 minutes resulted in the dramatic downregulation of claudin-5 expression at the cell junctions. Expression of VE-cadherin was not affected although it assumed a discontinuous (zigzag) pattern. Treatment with the ROCK inhibitor Y-27632 for 6 hours resulted in the change in the morphology of HLECs although expression of VE-Cadherin and claudin-5 were unaffected. Pre-treatment with Y-27632 inhibited the W146-induced downregulation of claudin-5. *(B, C) Inhibition of S1PR1 signaling in HLECs results in the activation of RhoA*. HLECs were (B) treated with W146 for the indicated time points or (C) transfected with siS1PR1 and RhoA activity was measured using the protein lysates. RhoA activity was enhanced rapidly by both W146 and siS1PR1. Statistics: n=3 for all experiments. *p < 0.05, **p < 0.01. Error bars in graphs represent ±SEM.

We next investigated how S1PR1 regulates Claudin-5 localization. Cell-cell interactions and cell shape are regulated by the actin cytoskeleton. Hence, we analyzed the cytoskeletal architecture of control and W146-treated HLECs by using Alexa 488-conjugated phalloidin staining. Cortical actin (actin around the cell periphery) was enriched in vehicle-treated HLECs (**Supplementary Figure 5A**). In contrast, treatment with W146 for 30 minutes induced active remodeling of cytoskeleton with the formation of radial actin bundles that are orientated perpendicularly to the cell periphery (**Supplementary Figure 5A**, red arrowheads). The increased accumulation of the radial actin fibers supports a cell retraction mechanism, which was also supported by the decreased size and compromised barrier integrity of W146-treated HLECs (**Supplementary Figure 5A** and **Supplementary Figure 6**). Altogether, these data show that W146 treatment compromises cytoskeletal architecture in cultured lymphatic endothelial cells.

The Rho family of small GTPases, such as RhoA, regulate cytoskeletal dynamics (47). We found that RhoA activity was increased in HLECs treated with W146 (**Figure 6B**). We tested whether inhibition of RhoA/ROCK pathway could restore Claudin-5 localization in W146-treated HLECs. Treatment of HLECs with ROCK inhibitor Y-27632 for 6 hours resulted in “oak leaf shaped” HLECs in which the expression of VE-Cadherin and Claudin-5 were intact (**Figure 6A**). Strikingly, Y-27632 pretreatment followed by incubation with W146 for 30 minutes prevented the formation of radial actin bundles, cellular retraction and Claudin-5 downregulation (**Figure 6A**). These results indicate that S1PR1 regulates the membrane localization of Claudin-5 in HLECs by inhibiting RhoA/ROCK activity. Nevertheless, HLEC permeability was increased by Y-27632 as well (**Supplementary Figure 6**). Furthermore, W146-induced formation of gaps between HLECs was not completely abolished by Y-27632 (**Supplementary Figure 5A**, white arrowheads and **Supplementary Figure 6**). These results indicate that both over-activation or inhibition of RhoA activity could compromise HLEC barrier integrity.

Localization of Claudin-5 at the periphery of HLECs was restored 24 hours after treatment with W146 (**Supplementary Figure 5B**). VE-Cadherin was also expressed in a uniform manner around the cell membrane, indicating that the cell-cell junctions were fully restored. However, treatment with W146 for 24 hours resulted in elongated HLECs with obvious increases in stress fiber (actin fibers that traverse across the cytoplasm) formation (**Supplementary Figure 5C**). The elongated morphology of HLECs, increased formation of stress fibers and elevated RhoA activity were also observed in HLECs treated with *siS1PR1* (**Figure 6C** and **Supplementary Figure 5D**). Based on these observations, we conclude that loss of S1PR1 upregulates RhoA activity and transiently disrupt Claudin-5 localization and cell-cell interactions. As Claudin-5 expression is unaffected in *siS1PR1*-treated HLECs, we concluded that Claudin-5 is dispensable for the crosstalk between LSS and VEGF-C signaling.

In conclusion, LSS regulates lymphangiogenesis via two distinct mechanisms of action. First, LSS inhibits the expression of pro-lymphangiogenic sprouting genes, such as DLL4, ANGPT2, ESM1 and ADM, which does not involve S1PR1. Second, LSS enhances VEGF-C, a process that is directly antagonized by S1PR1. Additionally, S1PR1 regulates the membrane localization of Claudin-5 by inhibiting RhoA activity. Correspondingly, *S1pr1* deletion in LECs results in hypersprouting and hyperbranching of lymphatic vessels, which are immature due to the delayed expression of Claudin-5 and abnormal localization of VE-Cadherin. Our findings are schematically summarized in **Figure 7**.

**Figure 7:**
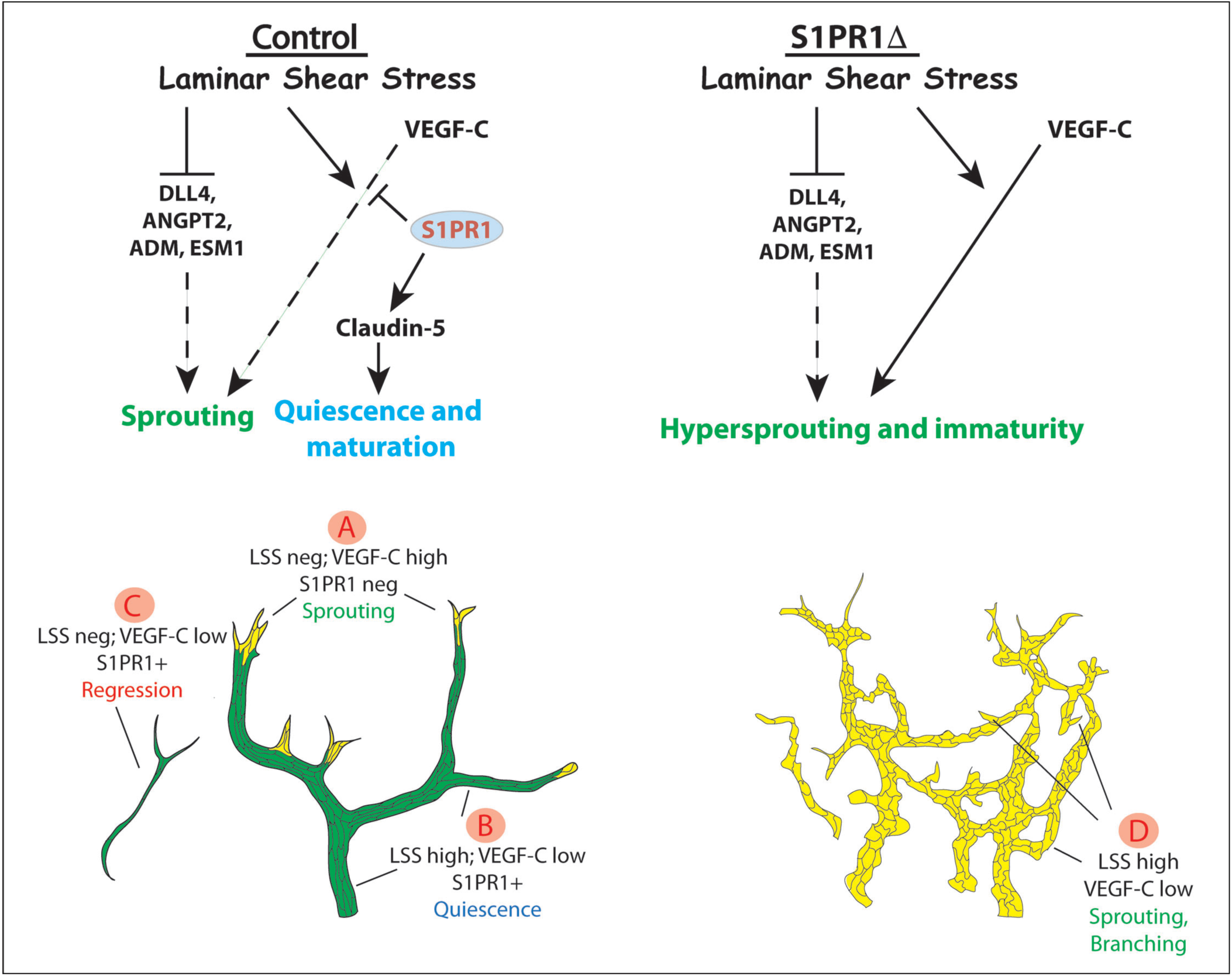
Model for the crosstalk between LSS, S1PR1 and VEGF-C signaling during lymphatic vascular patterning. A) LSS inhibits the expression of molecules such as DLL4, ANGPT2, ADM and ESM1 thereby limiting their expression to tip cells. B) LSS enhances VEGF-C signaling to promote the survival of quiescent LECs. S1PR1 functions as a buffer that prevents the hyper activation of LSS/VEGF-C signaling to prevent the sprouting of lymphatic vessels. C) Lymphatic vessels that are exposed to low dose of VEGF-C with little or no lymph flow regress due to reduced LSS/VEGF-C signaling. (D) In the absence of S1PR1, VEGF-C signaling is hyperactive in the posterior lymphatic plexus, which results in excessive number of sprouts and branches. In addition to modulating LSS/VEGF-C signaling, S1PR1 inhibits RhoA activity to regulate the cytoskeleton and maintain claudin-5 expression (green vessels). Consequently, mice lacking S1PR1 have immature cell junctions (yellow vessels).

## DISCUSSION

We have determined that the lymphatic vasculature-specific roles of DLL4 and S1PR1 are distinct from their known activities in blood endothelial cells. DLL4 is known to inhibit angiogenesis. In contrast, we have determined that DLL4 promotes lymphangiogenesis. S1PR1 is known to regulate the expression of VE-Cadherin and the alignment of blood endothelial cells with respect to LSS. In contrast, S1PR1 is not required for the expression of VE-Cadherin in HLECs or for the alignment of HLECs with respect to LSS. However, S1PR1 is required for the expression of claudin-5 in LECs in vitro and in vivo. Finally, while S1PR1 antagonizes VEGF-A signaling in blood vascular endothelial cells, it inhibits VEGF-C signaling that is specifically enhanced by LSS.

We have also identified several previously unknown mechanisms that operate during lymphatic vascular patterning. The tip-cell expressed molecule DLL4 is an enhancer of VEGF-C signaling. *Dll4*^*+/-*^ embryos have hypoplastic lymphatic vessels, establishing the pro-lymphangiogenic role of this gene. We currently do not know whether the other tip-cell expressed molecules ADM, ANGPT2 or ESM1 could regulate VEGF-C signaling. However, evidence from published manuscripts suggests that they might. Conditional deletion of *Calcrl*, the gene encoding the ADM receptor, in LECs results in a lacteal phenotype similar to that of *Dll4*^*+/-*^ mice (48). Furthermore, *CALCRL* knockdown downregulates, while ADM treatment enhances, *DLL4* expression in cultured HLECs (48). Therefore, ADM could regulate VEGF-C signaling indirectly through DLL4. ESM1 can enhance VEGF-A signaling in blood endothelial cells (6). In addition, ESM1 is able to enhance VEGF-C-mediated HLEC proliferation (7). Finally, ectopic expression of ANGPT2 promotes lymphangiogenesis, which could be inhibited by a VEGFR3-blocking antibody (49). Thus, DLL4, ANGPT2, ADM and ESM1 are likely part of a network that enhances VEGF-C signaling in lymphatic tip cells.

LSS is a potent inhibitor of *DLL4, ADM, ESM1* and *ANGPT2* expression in HLECs. Previous reports have shown that VEGF-C could transiently enhance *DLL4* expression in HLECs (4). Thus, a balance between VEGF-C and LSS activities might confine the expression of DLL4 and other molecules to non-perfused tip cells (**Figure 7A**). The shear-responsive transcription factor KLF2 is a potential candidate through which LSS inhibits the expression of tip cell genes, as KLF2 is known to inhibit *ANGPT2* and *EGR1* expressions in blood vascular endothelial cells (50).

We have determined that LSS enhances VEGF-C signaling in HLECs. Previous reports, including our own, have shown that *Clec2*^*-/-*^ mice, which lack lymph flow (51), have hypoplastic lymphatic vessels (13). Therefore, the LSS/VEGF-C signaling pathway is likely to be physiologically significant to promote the growth of lymphatic vessels.

LSS and MAPK signaling pathway are critical for the survival of quiescent blood vascular endothelial cells (44, 45, 52, 53). MAPK signaling is activated by FGF to maintain the survival and integrity of quiescent blood vascular endothelial cells (52). The mechanisms that activate MAPK signaling in quiescent LECs are currently unknown. However, deletion of *Vegfc* from adult mice results in the regression of lymphatic vessels (54). Therefore, we suggest that LSS/VEGF-C signaling could be important for the survival of quiescent LECs (**Figure 7B**). Coincidently, FGFR3 is one of the molecules that are upregulated by LSS in HLECs (19). Whether LSS could enhance FGF or other receptor tyrosine kinase signaling pathways remains to be determined.

In addition to promoting the growth and quiescence of LECs, LSS/VEGF-C signaling could eliminate redundant vessel segments that do not support physiologically productive lymph flow (**Figure 7C**). Accordingly, poorly perfused lymphatic vessels will experience reduced VEGF-C/VEGFR3 signaling compared to vessels with stronger lymph flow, and be pruned. Successive rounds of pruning will progressively strengthen lymph flow in the remaining vessels and result in their quiescence and maturation.

S1PR1 activity is excluded from tip cells. In contrast, S1PR1 activity is enriched in more-mature lymphatic vessels where it antagonizes LSS/VEGF-C signaling. S1PR1 activity is likely regulated by S1P that is present at high concentrations in lymph (28). We postulated that S1PR1 functions as a buffer that prevents the over activation of the LSS/VEGF-C signaling. This model would predict excessive sprouting and insufficient pruning of lymphatic vessels in *Lyve1-Cre;S1pr1*^*-/f*^ mice, resulting in excessive lymphatic branching, as we observed (**Figure 7D**).

Oscillatory shear stress is thought to be necessary for the maturation of lymphatic vessels (51). Approaches to determine lymph flow during development currently do not exist. The elongated morphology of LECs in embryonic lymphatic vessels suggests that they are likely exposed to LSS (**Figure 2A**). Nevertheless, LECs could be exposed to oscillatory shear stress at earlier developmental time points, and at branch points and valves (16). Whether oscillatory shear stress could enhance VEGF-C signaling at such instances remains to be determined.

S1PR1 plays an additional role in regulating Claudin-5 localization to LEC-junctions in developing lymphatic vessels. Claudin-5 expression is defective in the lymphatic vessels of *Lyve1-Cre;S1pr1*^*-/f*^ mice. S1PR1-mediated inhibition of RhoA signaling appears to be important for the localization of Claudin-5 to cell membrane. Intriguingly, RhoA activity also appears to be important for LEC junctional integrity, as inhibition of ROCK results in abnormally shaped HLECs.

Due to the similarities between the blood and lymphatic endothelium, we are tempted to suggest that our findings could be relevant during developmental and pathological angiogenesis as well. Whether S1PR1 function modulating drugs could be used to treat angiogenic defects should be tested. Furthermore, T-cell expressed S1PR1 is an important target for treating the neuroinflammatory disease multiple sclerosis (55). Recent evidences have suggested that meningeal lymphatic vessels could regulate neuroinflammation (56). It will be important to determine how FDA approved drugs such as Fingolimod could affect the structure and function of meningeal lymphatic vessels.

## MATERIALS AND METHODS

IHC on sections and skin were performed according to our previously published protocols (13, 14, 57). Immunocytochemistry was performed as we described recently (13, 25).

### Antibodies

Primary antibodies for immunohistochemistry: rabbit anti-PROX1 (Cat# 11-002, Angiobio, San Diego, CA, USA), goat anti-human PROX1 (Cat# AF2727, R&D Systems, Minneapolis, MN, USA), goat anti-mouse VEGRF3 (Cat# AF743, R&D Systems, Minneapolis, MN, USA), rat anti-mouse S1PR1 (Cat# MAB7089, R&D Systems, Minneapolis, MN, USA), rat anti-mouse CD31 (Cat# 553370, BD Pharmingen, San Jose, CA, USA), rat anti-mouse VE-Cadherin (Cat# 550548, BD Pharmingen, San Jose, CA, USA), chicken anti-GFP (Cat# ab13970, abcam, Cambridge, MA, USA), goat anti-mouse LYVE-1 (Cat# AF2125, R&D Systems, Minneapolis, MN, USA), rabbit anti-human ZO-1 (Cat# 40-2200, Invitrogen, Camarillo, CA, USA), rabbit anti-mouse CLDN5 (Cat #34-1600, Thermo Fisher Scientific, Rockford, IL, USA) and Alexa 488-conjugated Phalloidin (Cat# A12379, Invitrogen, Camarillo, CA, USA).

Secondary antibodies for immunohistochemistry: Cy3-conjugated donkey anti-rabbit, Cy3-conjugated donkey anti-Rat, Cy3-conjugated donkey anti-goat, Cy5-conjugated donkey anti-rat, Cy5-conjugated donkey anti-goat, FITC-conjugated donkey anti-chicken and Biontin-conjugated donkey anti-goat antibodies were purchased from Jackson ImmunoResearch Laboratories (West Grove, PA, USA). Alexa 488-conjugated donkey anti-goat and Alexa 488-conjugated donkey anti-rat antibodies were purchased from Life Technologies (Grand Island, NY, USA).

Primary antibodies for Western blotting: mouse anti-β-Actin (Cat# A5441, Sigma-Aldrich, St. Louis, MO, USA), rabbit anti-human GAPDH (Cat# PAB13195, Abnova, Taipei City, Taiwan, R.O.C), rabbit anti-human S1PR1 (Cat#A12935, ABclonal, Woburn, MA) and rabbit anti-human eNOS (Cat#NB300-500SS, Novus Biologicals). Rabbit anti-mouse AKT (Cat# 4691), rabbit anti-human Phospho-AKT (Cat# 4060), rabbit anti-rat ERK1/2 (Cat# 4695), rabbit anti-human p38 MAPK (Cat# 8690), rabbit anti-human Phospho-p38 MAPK (Cat# 4511), rabbit anti-human Phospho-eNOS (Cat# 9571), rabbit anti-human NICD (Cat# 4147), rabbit anti-human DLL4 (Cat# 2589) and rabbit anti-RhoA (Cat# 2117) were purchased from Cell Signaling Technology, Danvers, MA.

HRP-conjugated secondary antibodies for Western blotting: goat anti-mouse IgG (Cat# A 4416, Sigma-Aldrich, St. Louis, MO, USA) and goat anti-rabbit IgG (Cat# GtxRb-003-EHRPX, ImmunoReagents, Raleigh, NC).

### Cells

Dr. Donwong Choi provided the HLECs harvested according to their reported protocols (15, 18, 19). HLECs were grown on culture dishes or glass slide coated with 0.2% gelatin and were maintained in EGM-2 EC Growth Medium-2 Bullet Kit (Lonza). All experiments were conducted using cells until passage (P) 8. HLECs were treated as potential biohazards and were handled according to institutional biosafety regulations.

### Cell treatments

#### S1PR1 agonist and antagonist treatment

HLECs were seeded onto six-well plates. After 48 hr culture (80 to 100% confluency), cells were starved for six hours and then treated with 20 μM SEW2871 (Cat# 10006440, Cayman Chemical, Ann Arbor, MI), 10 μM W146 (Cat# 10009109, Cayman Chemical, Ann Arbor, MI) or vehicle respectively for one hour. Pretreated HLECs were subsequently treated with 100ng/ml VEGF-C (cat# 9199-vc-025/CF, R&D Systems) for 10 minutes.

#### siRNA transfection

HLECs were seeded onto six-well plates or culture dishes. After 24 hours of culture (around 30% confluency), cells were transfected with siCTR (Cat# 51-01-14-03, Integrated DNA Technologies), siS1PR1 (Cat# SI00376201, Qiagen), or siDLL4 (hs.Ri.DLL4.13.1, Integrated DNA Technologies) using Lipofectamine RNAiMax Tranfection Reagent (Cat# 13778150, Invitrogen) according to manufacturer’s instruction.

#### VEGF treatment

Confluent HLECs were treated with the indicated concentrations of VEGF-C (cat# 9199-vc-025/CF, R&D Systems) or VEGF-A (cat# CYT-241, ProSpec) and collected after the indicated time points.

#### Laminar Shear Stress

HLECs were seeded in culture dishes and grown to 80-100% confluency before exposing them to LSS for either 10 minutes or 24 hours. LSS was applied to the cells at 5 dynes/cm^2^ using the approach described previously (18, 19). Cells were not serum starved for VEGF-C treatment after LSS. VEGF-C was added to the dishes and incubated for 10-30 minutes for Western blotting. For quantifying *EGR1* expression VEGF-C was added to the cells and returned back to LSS for 1 hour.

#### Rock inhibitor treatment

HLECs were seeded onto 24-well plates. 100% confluent cells were treated with 10 μM Y-27632 (Cat# SCM075, Sigma) in starvation media for six hours. The cells were then treated with either vehicle or 10 μM W146 for 30 minutes. Control cells were grown in starvation media for 6 hours followed by vehicle or 10 μM W146 for 30 minutes.

### Cell junction analysis

HLECs were treated with chemicals as described above and their intracellular permeability was analyzed using In Vitro Vascular Permeability Imaging Assay (Cat# 17-10398, Sigma) according to manufacturer’s instructions.

### Mice

*Lyve1-Cre* (28), *S1pr1*^*flox*^ (29), *S1PR1-GFP* (26) mice were described previously and were purchased from Jackson laboratory (catalog numbers 012601, 019141 and 026275 respectively). Mice were maintained in C57BL6 or C57BL6/NMRI mixed backgrounds. *Dll4*^*+/-*^ cryopreserved embryos were purchased from the Canadian Mouse Mutant Repository (CMMR) and implanted into CD-1/ICR females then backcrossed onto a C57BL6 background (34, 58). HI provided the *Vegfr3*^*+/EGFP*^ mice (59).

### Protein isolation and analysis

Protein was extracted from cells by using RIPA lysis buffer. Western blots were performed according to standard protocols. The protein lysate was not boiled while performing Western blot for S1PR1.

### RNA isolation and quantitative real-time PCR

Total RNA from HLECs was purified using Trizol (Invitrogen, Carlsbad, CA, USA) according to manufacturers instructions. cDNA was synthesized from total RNA (0.1 – 1.0 µg) with iScript Advanced cDNA Synthesis Kit (BioRad, Hercules, CA, USA). QRT-PCR was performed using PowerUp SYBR Green Master Mix reagent (Applied Biosystems, CA, USA) in a CFX96 Real-Time System (Bio-Rad, Hercules, CA, USA). Expression levels were normalized to GAPDH. Primer sequences will be provided upon request.

### Rho GTPase pull-down assays

For RhoA pull-down experiments we used the GST-RBD plasmid, a gift from Dr. Martin Schwartz (Addgene plasmid # 15247; http://n2t.net/addgene:15247; RRID:Addgene_15247). RhoA activation in cultured cells was assessed as previously described (60). Briefly, after treatment, the cells were lysed in lysis buffer containing 20mM Hepes, pH 7.4, 0.1M NaCl, 1% Triton X-100, 10mM EGTA, 40mM b-glycerophosphate, 20mM MgCl_2_, 1mM Na_3_VO_4_, 1mM dithiothreitol, 10 µg/ml aprotinin, 10 µg/ml leupeptin and 1mM phenylmethylsulfonyl fluoride on ice. The lysates were incubated with the glutathione S-transferase-rhotekin-Rho-binding domain previously bound to glutathione-Sepharose beads (Amersham Biosciences) and washed three times with lysis buffer. Associated GTP-bound forms of Rho were released with SDS–polyacrylamide gel electrophoresis loading buffer and analyzed by Western blot analysis using a monoclonal antibody against RhoA (Cat #2117, Cell Signaling). Normalization took place based on the total RhoA levels present in the whole cell lysate of the same samples.

### Statistics

For biochemical studies the number n refers to the number of times the experiment was performed. For histochemical analysis n refers to the total number of animals included per group. Statistically significant differences were determined using unpaired t test. Prism software was used for statistical analyses. Data are reported as mean ± SEM with significance set at P < 0.05. n and p values for each experiment is provided in the figure legends. Western blots and qRT-PCR’s are performed at least three independent times. The most representative Western blots are presented. The average values from individual qRT-PCR assays were presented unless otherwise stated.

### Study Approval

All mice were housed and handled according to the institutional IACUC protocols.

## AUTHOR CONTRIBUTIONS

XG, TH and RSS conceptualized the work; XG, KY, RA, LC, YCH, KBR, CM, and RSS performed experiments; DC, HC, JDW, HI and TH provided critical reagents; XG and RSS wrote the manuscript with critical input from JDW; all authors provided input in designing the experiments and in editing the manuscript.

## ACKNOWLEDGEMENTS

This work is supported by NIH/NHLBI (R01HL131652 to RSS; R01HL133216 to RSS and HC, R35HL135821 to TH), NIH/NIGMS COBRE (P20 GM103441 to XG; PI: Dr. McEver), Oklahoma Center for Adult Stem Cell Research (4340) to RSS and American Heart Association (19POST34380819 to YH). We thank Drs. Wayne Orr and Martin Schwartz for helpful suggestions and reagents and Mr. Judson Copeland for the illustration. We thank Dr. Angela Andersen, Life Science Editors, for editing the manuscript.

## Supplementary Figures and Figure legends

**Supplementary Figure 1:**
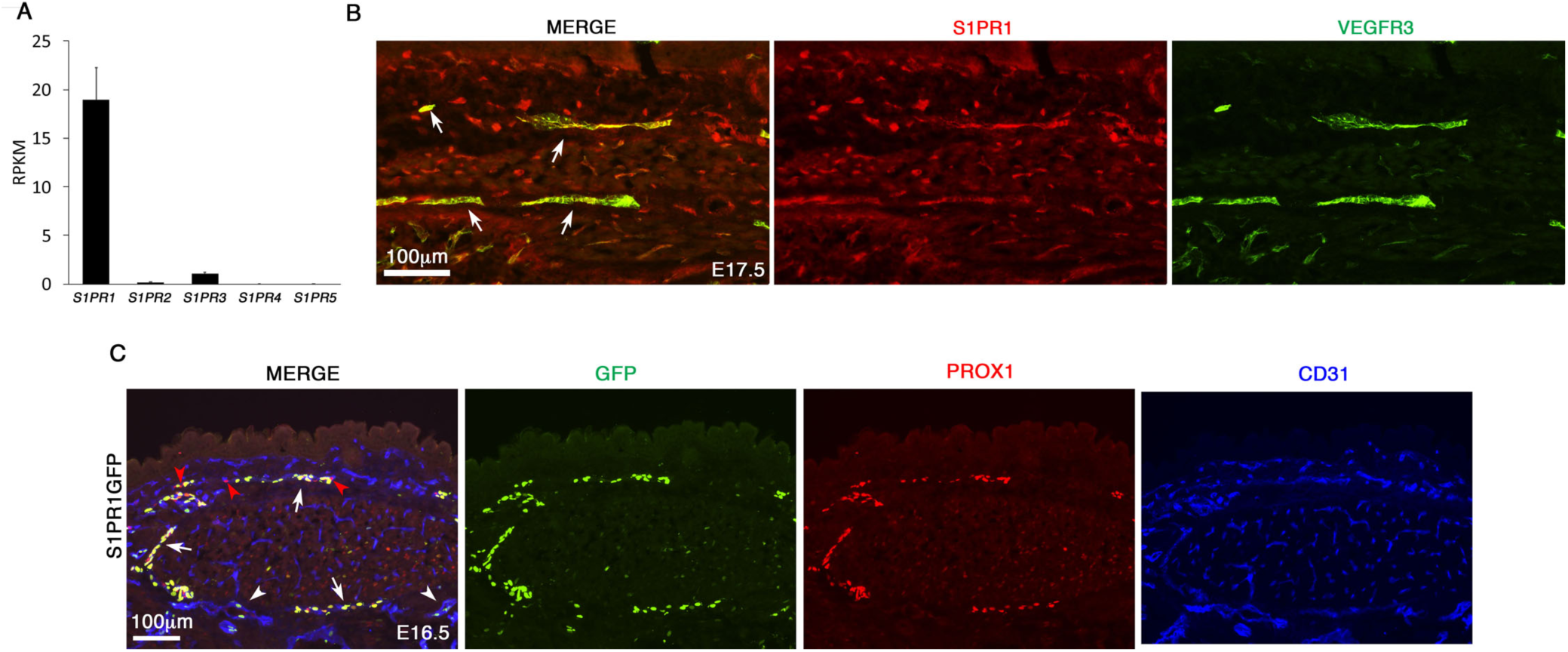
S1PR1 expression and activity are observed in HLECs and mouse LECs. *(A) S1PR1 is the most strongly expressed S1P receptor in HLECs*. RNA-seq was performed using primary human LECs and the reads per kilo base of transcript per million mapped reads (RPKM) for the five S1P receptors were plotted. The RPKM values suggest that S1P1 is the most enriched S1P receptor in HLECs. *(B) S1PR1 is expressed in developing murine lymphatic vessels*. E17.5 embryos were sectioned and analyzed by immunohistochemistry. S1PR1 was expressed in the VEGFR3^+^ lymphatic vessels of the embryos. Statistics: n=4. *(C) Heterogeneous S1PR1 activity is observed in murine LECs*. S1PR1-GFP embryos were sectioned and immunostained for GFP. A few GFP^+^ blood endothelial cells were observed (arrowheads). Most LECs were GFP^+^ (arrows) although a few GFP^-^ LECs were also observed (red arrowheads). Statistics: n=4 for both S1PR1GFP and H2B-GFP.

**Supplementary Figure 2:**
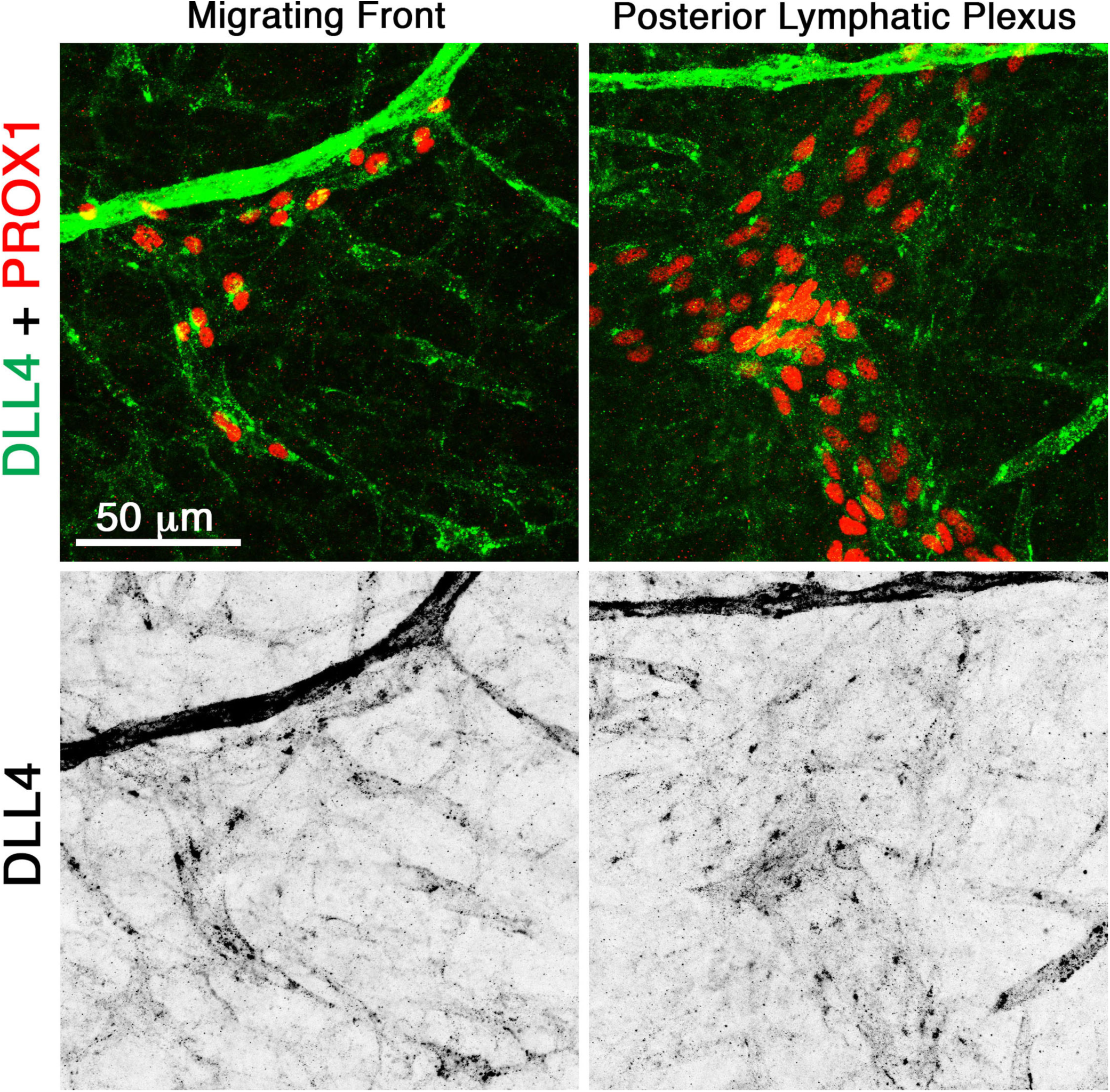
DLL4 expression is enriched in the migrating front of lymphatic vessels. Dorsal skins of E16.5 wild type embryos were analyzed for the expression of DLL4. DLL4 was expressed in the migrating front of lymphatic vessels. In contrast, DLL4 expression in the more-mature and quiescent lymphatic vessels was greatly reduced. Statistics: n=5.

**Supplementary Figure 3:**
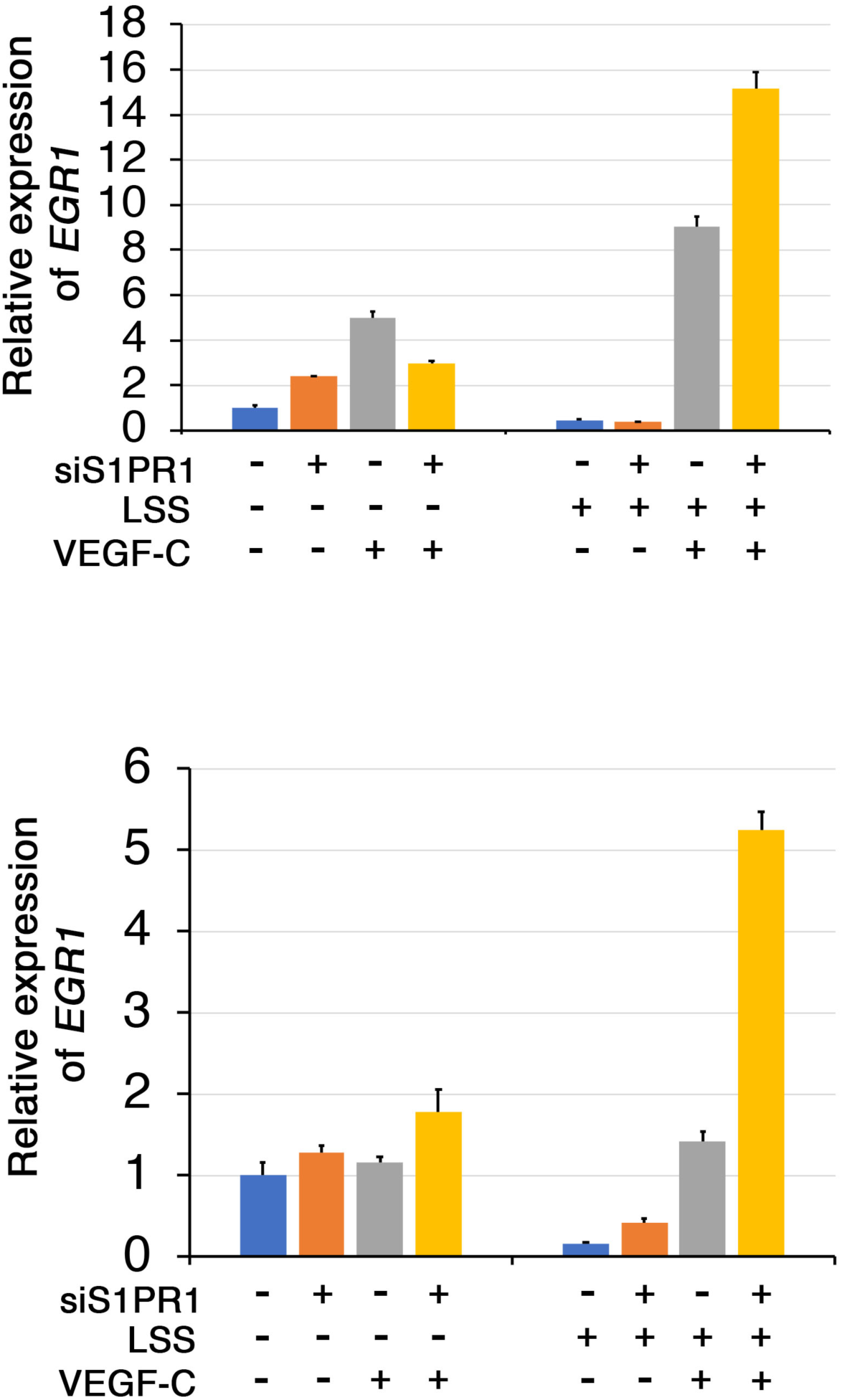
S1PR1 antagonizes LSS/VEGF-C signaling induced expression of EGR1. VEGF-C induced expression of *EGR1* was dramatically enhanced by LSS and it was further enhanced by siS1PR1. Statistics: Data from 2 out of 3 independent experiments is presented here. Data from the other experiment is presented in Figure 5C. P values were not presented due to large variability in fold activation of *EGR1* between independent experiments

**Supplementary Figure 4:**
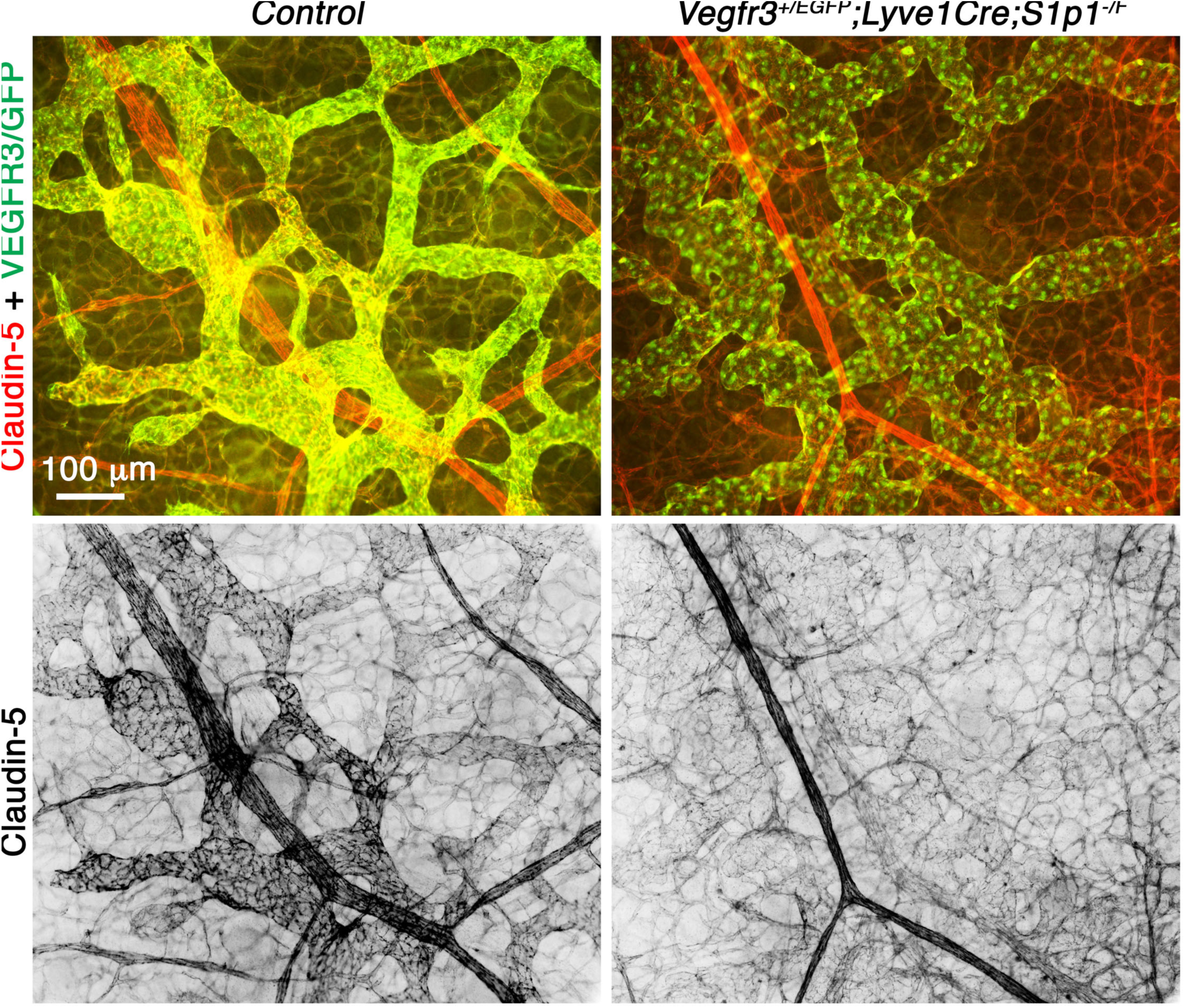
Heterozygosity of *Vegfr3* could not rescue the expression of claudin-5 in embryos lacking S1PR1. Claudin-5 was expressed in a gradient manner within the growing lymphatic vessels of control embryos with weaker expression in the migrating tips and stronger expression in the vessels behind. Claudin-5 expression was uniformly downregulated in the lymphatic vessels of *Lyve1-Cre;S1r1*^*-/f*^ embryos lacking one allele of *Vegfr3*. Statistics: n=5 for control embryos and n=4 for mutant embryos.

**Supplementary Figure 5:**
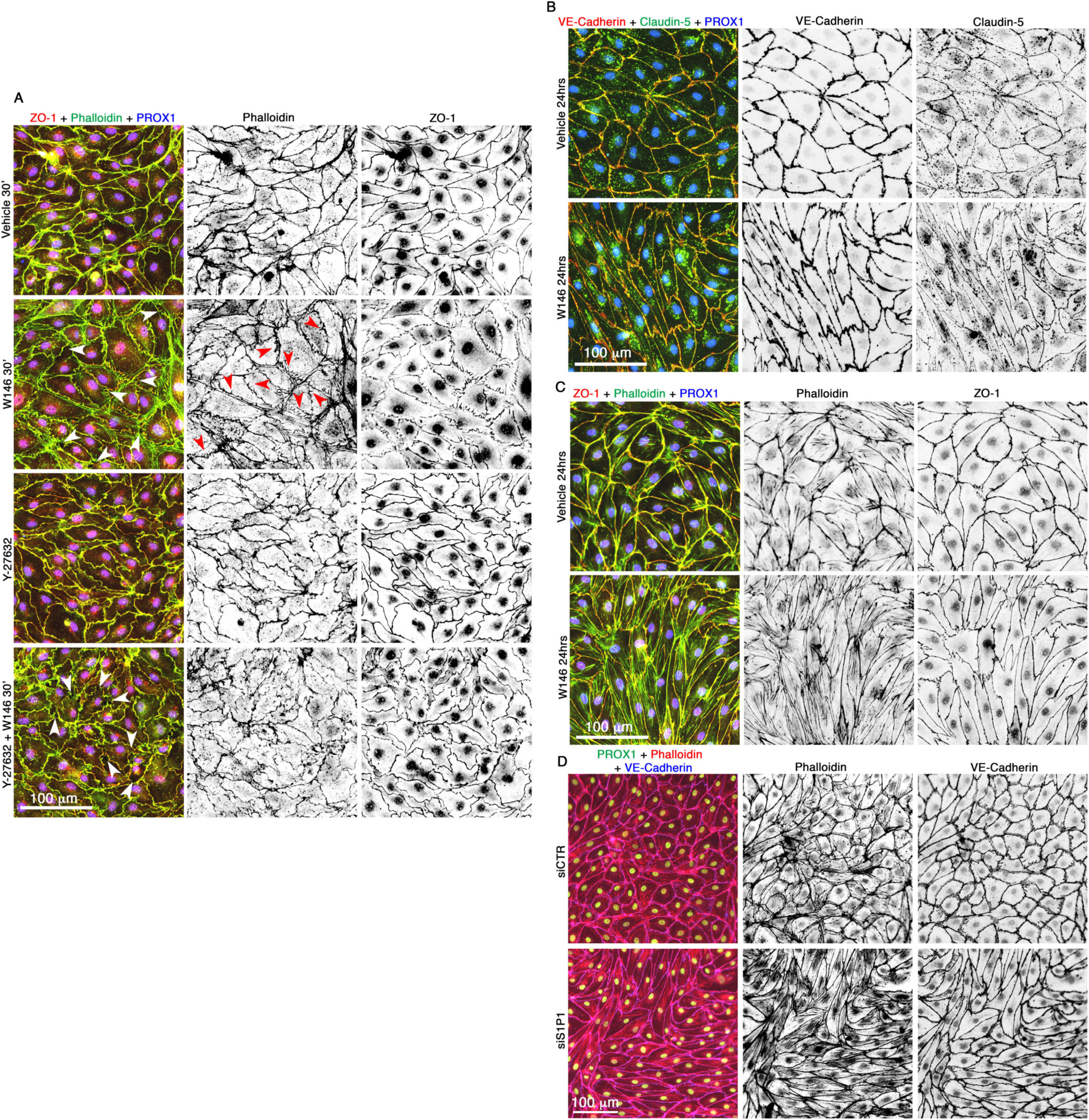
Inhibition of S1PR1 promotes cell junctional and cytoskeletal defects in HLECs. *(A) S1PR1 antagonist W146 triggers cytoskeletal reorganization in HLECs, which could be prevented by the inhibition of ROCK*. Phalloidin staining revealed the presence of actin filaments along the periphery of cells (cortical actin) in vehicle treated HLECs. Tight junction molecule ZO-1 was uniformly expressed along the cell membrane. Treatment of HLECs with W146 for 30 minutes resulted in the formation of radial actin bundles, which were aligned perpendicular to cell membrane (red arrowheads). W146 treatment also resulted in the discontinuous expression pattern of ZO-1. Treatment of HLECs with the ROCK inhibitor Y-227632 for 6 hours promoted cell shape change. However, Y-227632 treatment did not cause any obvious defects in the expression of actin or ZO-1. Pretreatment of HLECs with Y-227632 for 6 hours dramatically inhibited the formation of radial filaments upon W146 treatment. White arrowheads point to the intracellular gaps. *(B-D) Prolonged inhibition of S1PR1 results in the formation of stress fibers and cell elongation*. (B, C) Treatment with W146 for 24 hours resulted in elongated HLECs (B) in which stress fibers could be seen traversing the cytoplasm (C). No obvious defects were observed in the expressions of VE-Cadherin, claudin-5 or ZO-1. (D) Knockdown of S1PR1 from HLECs also resulted in the elongated cells with numerous stress fibers. Statistics: n=3 for all experiments.

**Supplementary Figure 6:**
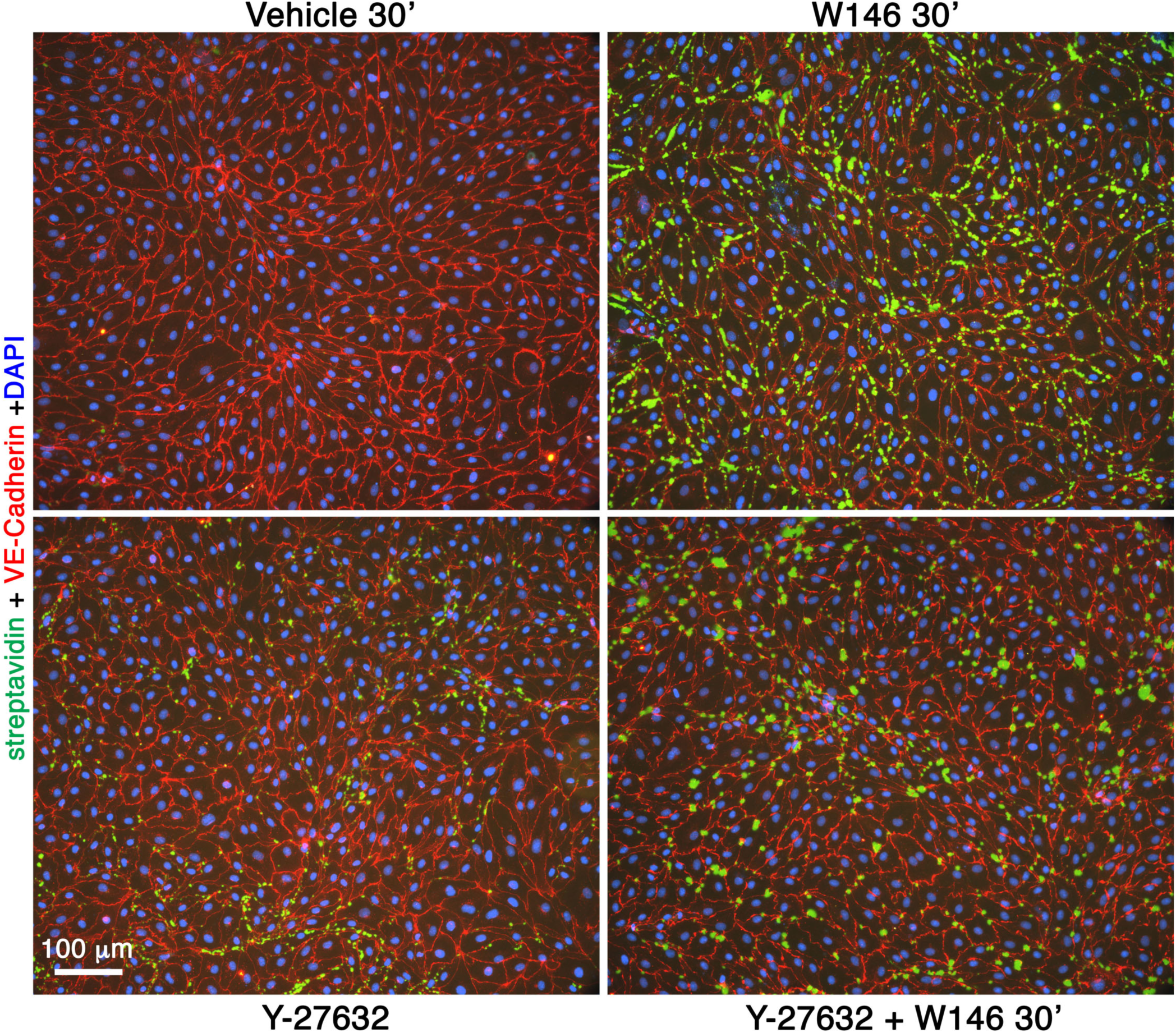
W146 increases the permeability of HLECs and it could not be ameliorated by the inhibition of ROCK. Confluent HLECs were treated with W146 for 30 minutes with or without pretreatment with ROCK inhibitor Y-227632 for 6 hours. Intracellular gaps (green signals) were increased by W146. Y-227632 also modestly increased the intracellular gaps. Y-227632 was also not able to prevent the formation of intracellular gaps by W146. Statistics: n=3.

## REFERENCES

1. Adams RH, and Alitalo K. Molecular regulation of angiogenesis and lymphangiogenesis. Nature reviews Molecular cell biology. 2007;8(6):464–78.

2. Gerhardt H, Golding M, Fruttiger M, Ruhrberg C, Lundkvist A, Abramsson A, et al. VEGF guides angiogenic sprouting utilizing endothelial tip cell filopodia. J Cell Biol. 2003;161(6):1163–77.

3. del Toro R, Prahst C, Mathivet T, Siegfried G, Kaminker JS, Larrivee B, et al. Identification and functional analysis of endothelial tip cell-enriched genes. Blood. 2010;116(19):4025–33.

4. Dieterich LC, Klein S, Mathelier A, Sliwa-Primorac A, Ma Q, Hong YK, et al. DeepCAGE Transcriptomics Reveal an Important Role of the Transcription Factor MAFB in the Lymphatic Endothelium. Cell reports. 2015;13(7):1493–504.

5. Lobov IB, Renard RA, Papadopoulos N, Gale NW, Thurston G, Yancopoulos GD, et al. Delta-like ligand 4 (Dll4) is induced by VEGF as a negative regulator of angiogenic sprouting. Proceedings of the National Academy of Sciences of the United States of America. 2007;104(9):3219–24.

6. Rocha SF, Schiller M, Jing D, Li H, Butz S, Vestweber D, et al. Esm1 modulates endothelial tip cell behavior and vascular permeability by enhancing VEGF bioavailability. Circulation research. 2014;115(6):581–90.

7. Shin JW, Huggenberger R, and Detmar M. Transcriptional profiling of VEGF-A and VEGF-C target genes in lymphatic endothelium reveals endothelial-specific molecule-1 as a novel mediator of lymphangiogenesis. Blood. 2008;112(6):2318–26.

8. Vaahtomeri K, Karaman S, Makinen T, and Alitalo K. Lymphangiogenesis guidance by paracrine and pericellular factors. Genes Dev. 2017;31(16):1615–34.

9. Karkkainen MJ, Haiko P, Sainio K, Partanen J, Taipale J, Petrova TV, et al. Vascular endothelial growth factor C is required for sprouting of the first lymphatic vessels from embryonic veins. Nat Immunol. 2004;5(1):74–80.

10. Karkkainen MJ, Saaristo A, Jussila L, Karila KA, Lawrence EC, Pajusola K, et al. A model for gene therapy of human hereditary lymphedema. Proceedings of the National Academy of Sciences of the United States of America. 2001;98(22):12677–82.

11. Yao LC, Testini C, Tvorogov D, Anisimov A, Vargas SO, Baluk P, et al. Pulmonary lymphangiectasia resulting from vascular endothelial growth factor-C overexpression during a critical period. Circulation research. 2014;114(5):806–22.

12. Baeyens N, Bandyopadhyay C, Coon BG, Yun S, and Schwartz MA. Endothelial fluid shear stress sensing in vascular health and disease. The Journal of clinical investigation. 2016;126(3):821–8.

13. Cha B, Geng X, Mahamud MR, Fu J, Mukherjee A, Kim Y, et al. Mechanotransduction activates canonical Wnt/beta-catenin signaling to promote lymphatic vascular patterning and the development of lymphatic and lymphovenous valves. Genes Dev. 2016;30(12):1454–69.

14. Cha B, Geng X, Mahamud MR, Zhang JY, Chen L, Kim W, et al. Complementary Wnt Sources Regulate Lymphatic Vascular Development via PROX1-Dependent Wnt/beta-Catenin Signaling. Cell reports. 2018;25(3):571–84 e5.

15. Choi D, Park E, Jung E, Cha B, Lee S, Yu J, et al. Piezo1 incorporates mechanical force signals to genetic program that governs lymphatic valve development and maintenance. JCI Insight. 2019.

16. Sabine A, Agalarov Y, Maby-El Hajjami H, Jaquet M, Hagerling R, Pollmann C, et al. Mechanotransduction, PROX1, and FOXC2 cooperate to control connexin37 and calcineurin during lymphatic-valve formation. Developmental cell. 2012;22(2):430–45.

17. Wang Y, Baeyens N, Corti F, Tanaka K, Fang JS, Zhang J, et al. Syndecan 4 controls lymphatic vasculature remodeling during mouse embryonic development. Development. 2016;143(23):4441–51.

18. Choi D, Park E, Jung E, Seong YJ, Hong M, Lee S, et al. ORAI1 Activates Proliferation of Lymphatic Endothelial Cells in Response to Laminar Flow Through Kruppel-Like Factors 2 and 4. Circulation research. 2017;120(9):1426–39.

19. Choi D, Park E, Jung E, Seong YJ, Yoo J, Lee E, et al. Laminar flow downregulates Notch activity to promote lymphatic sprouting. The Journal of clinical investigation. 2017;127(4):1225–40.

20. Gaengel K, Niaudet C, Hagikura K, Lavina B, Muhl L, Hofmann JJ, et al. The sphingosine-1-phosphate receptor S1PR1 restricts sprouting angiogenesis by regulating the interplay between VE-cadherin and VEGFR2. Developmental cell. 2012;23(3):587–99.

21. Lee MJ, Thangada S, Claffey KP, Ancellin N, Liu CH, Kluk M, et al. Vascular endothelial cell adherens junction assembly and morphogenesis induced by sphingosine-1-phosphate. Cell. 1999;99(3):301–12.

22. Yanagida K, Engelbrecht E, Niaudet C, Jung B, Gaengel K, Holton K, et al. Sphingosine 1-Phosphate Receptor Signaling Establishes AP-1 Gradients to Allow for Retinal Endothelial Cell Specialization. Developmental cell. 2020.

23. Jung B, Obinata H, Galvani S, Mendelson K, Ding BS, Skoura A, et al. Flow-regulated endothelial S1P receptor-1 signaling sustains vascular development. Developmental cell. 2012;23(3):600–10.

24. Chien S. Mechanotransduction and endothelial cell homeostasis: the wisdom of the cell. Am J Physiol Heart Circ Physiol. 2007;292(3):H1209–24.

25. Mahamud MR, Geng X, Ho YC, Cha B, Kim Y, Ma J, et al. GATA2 controls lymphatic endothelial cell junctional integrity and lymphovenous valve morphogenesis through miR-126. Development. 2019.

26. Kono M, Tucker AE, Tran J, Bergner JB, Turner EM, and Proia RL. Sphingosine-1-phosphate receptor 1 reporter mice reveal receptor activation sites in vivo. The Journal of clinical investigation. 2014;124(5):2076–86.

27. Coxam B, Sabine A, Bower NI, Smith KA, Pichol-Thievend C, Skoczylas R, et al. Pkd1 regulates lymphatic vascular morphogenesis during development. Cell reports. 2014;7(3):623–33.

28. Pham TH, Baluk P, Xu Y, Grigorova I, Bankovich AJ, Pappu R, et al. Lymphatic endothelial cell sphingosine kinase activity is required for lymphocyte egress and lymphatic patterning. The Journal of experimental medicine. 2010;207(1):17–27.

29. Allende ML, Yamashita T, and Proia RL. G-protein-coupled receptor S1P1 acts within endothelial cells to regulate vascular maturation. Blood. 2003;102(10):3665–7.

30. Tzima E, Irani-Tehrani M, Kiosses WB, Dejana E, Schultz DA, Engelhardt B, et al. A mechanosensory complex that mediates the endothelial cell response to fluid shear stress. Nature. 2005;437(7057):426–31.

31. Hagerling R, Hoppe E, Dierkes C, Stehling M, Makinen T, Butz S, et al. Distinct roles of VE-cadherin for development and maintenance of specific lymph vessel beds. The EMBO journal. 2018;37(22).

32. Yang Y, Cha B, Motawe ZY, Srinivasan RS, and Scallan JP. VE-Cadherin Is Required for Lymphatic Valve Formation and Maintenance. Cell reports. 2019;28(9):2397–412 e4.

33. Yanagida K, Liu CH, Faraco G, Galvani S, Smith HK, Burg N, et al. Size-selective opening of the blood-brain barrier by targeting endothelial sphingosine 1-phosphate receptor 1. Proceedings of the National Academy of Sciences of the United States of America. 2017;114(17):4531–6.

34. Duarte A, Hirashima M, Benedito R, Trindade A, Diniz P, Bekman E, et al. Dosage-sensitive requirement for mouse Dll4 in artery development. Genes Dev. 2004;18(20):2474–8.

35. Hellstrom M, Phng LK, Hofmann JJ, Wallgard E, Coultas L, Lindblom P, et al. Dll4 signalling through Notch1 regulates formation of tip cells during angiogenesis. Nature. 2007;445(7129):776–80.

36. Gale NW, Dominguez MG, Noguera I, Pan L, Hughes V, Valenzuela DM, et al. Haploinsufficiency of delta-like 4 ligand results in embryonic lethality due to major defects in arterial and vascular development. Proceedings of the National Academy of Sciences of the United States of America. 2004;101(45):15949–54.

37. Krebs LT, Shutter JR, Tanigaki K, Honjo T, Stark KL, and Gridley T. Haploinsufficient lethality and formation of arteriovenous malformations in Notch pathway mutants. Genes Dev. 2004;18(20):2469–73.

38. Hogan BM, Herpers R, Witte M, Helotera H, Alitalo K, Duckers HJ, et al. Vegfc/Flt4 signalling is suppressed by Dll4 in developing zebrafish intersegmental arteries. Development. 2009;136(23):4001–9.

39. Bernier-Latmani J, Cisarovsky C, Demir CS, Bruand M, Jaquet M, Davanture S, et al. DLL4 promotes continuous adult intestinal lacteal regeneration and dietary fat transport. The Journal of clinical investigation. 2015;125(12):4572–86.

40. Zheng W, Tammela T, Yamamoto M, Anisimov A, Holopainen T, Kaijalainen S, et al. Notch restricts lymphatic vessel sprouting induced by vascular endothelial growth factor. Blood. 2011;118(4):1154–62.

41. Niessen K, Zhang G, Ridgway JB, Chen H, Kolumam G, Siebel CW, et al. The Notch1-Dll4 signaling pathway regulates mouse postnatal lymphatic development. Blood. 2011;118(7):1989–97.

42. Dellinger M, Hunter R, Bernas M, Gale N, Yancopoulos G, Erickson R, et al. Defective remodeling and maturation of the lymphatic vasculature in Angiopoietin-2 deficient mice. Developmental biology. 2008;319(2):309–20.

43. Fritz-Six KL, Dunworth WP, Li M, and Caron KM. Adrenomedullin signaling is necessary for murine lymphatic vascular development. The Journal of clinical investigation. 2008;118(1):40–50.

44. Akimoto S, Mitsumata M, Sasaguri T, and Yoshida Y. Laminar shear stress inhibits vascular endothelial cell proliferation by inducing cyclin-dependent kinase inhibitor p21(Sdi1/Cip1/Waf1). Circulation research. 2000;86(2):185–90.

45. Lin K, Hsu PP, Chen BP, Yuan S, Usami S, Shyy JY, et al. Molecular mechanism of endothelial growth arrest by laminar shear stress. Proceedings of the National Academy of Sciences of the United States of America. 2000;97(17):9385–9.

46. Baeyens N, Larrivee B, Ola R, Hayward-Piatkowskyi B, Dubrac A, Huang B, et al. Defective fluid shear stress mechanotransduction mediates hereditary hemorrhagic telangiectasia. J Cell Biol. 2016;214(7):807–16.

47. Kjoller L, and Hall A. Signaling to Rho GTPases. Experimental cell research. 1999;253(1):166–79.

48. Davis RB, Kechele DO, Blakeney ES, Pawlak JB, and Caron KM. Lymphatic deletion of calcitonin receptor-like receptor exacerbates intestinal inflammation. JCI Insight. 2017;2(6):e92465.

49. Zheng W, Nurmi H, Appak S, Sabine A, Bovay E, Korhonen EA, et al. Angiopoietin 2 regulates the transformation and integrity of lymphatic endothelial cell junctions. Genes Dev. 2014;28(14):1592–603.

50. Parmar KM, Larman HB, Dai G, Zhang Y, Wang ET, Moorthy SN, et al. Integration of flow-dependent endothelial phenotypes by Kruppel-like factor 2. The Journal of clinical investigation. 2006;116(1):49–58.

51. Sweet DT, Jimenez JM, Chang J, Hess PR, Mericko-Ishizuka P, Fu J, et al. Lymph flow regulates collecting lymphatic vessel maturation in vivo. The Journal of clinical investigation. 2015;125(8):2995–3007.

52. Murakami M, Nguyen LT, Zhuang ZW, Moodie KL, Carmeliet P, Stan RV, et al. The FGF system has a key role in regulating vascular integrity. The Journal of clinical investigation. 2008;118(10):3355–66.

53. Ricard N, Scott RP, Booth CJ, Velazquez H, Cilfone NA, Baylon JL, et al. Endothelial ERK1/2 signaling maintains integrity of the quiescent endothelium. The Journal of experimental medicine. 2019;216(8):1874–90.

54. Nurmi H, Saharinen P, Zarkada G, Zheng W, Robciuc MR, and Alitalo K. VEGF-C is required for intestinal lymphatic vessel maintenance and lipid absorption. EMBO Mol Med. 2015;7(11):1418–25.

55. Brinkmann V, Davis MD, Heise CE, Albert R, Cottens S, Hof R, et al. The immune modulator FTY720 targets sphingosine 1-phosphate receptors. J Biol Chem. 2002;277(24):21453–7.

56. Louveau A, Herz J, Alme MN, Salvador AF, Dong MQ, Viar KE, et al. CNS lymphatic drainage and neuroinflammation are regulated by meningeal lymphatic vasculature. Nat Neurosci. 2018;21(10):1380–91.

57. Geng X, Cha B, Mahamud MR, Lim KC, Silasi-Mansat R, Uddin MK, et al. Multiple mouse models of primary lymphedema exhibit distinct defects in lymphovenous valve development. Developmental biology. 2016;409(1):218–33.

58. Herman AM, Rhyner AM, Devine WP, Marrelli SP, Bruneau BG, and Wythe JD. A novel reporter allele for monitoring Dll4 expression within the embryonic and adult mouse. Biol Open. 2018;7(3).

59. Ichise T, Yoshida N, and Ichise H. H-, N- and Kras cooperatively regulate lymphatic vessel growth by modulating VEGFR3 expression in lymphatic endothelial cells in mice. Development. 2010;137(6):1003–13.

60. Zahra FT, Sajib MS, Ichiyama Y, Akwii RG, Tullar PE, Cobos C, et al. Endothelial RhoA GTPase is essential for in vitro endothelial functions but dispensable for physiological in vivo angiogenesis. Scientific reports. 2019;9(1):11666.

