## Supplementary Figures and Figure Legends for "S1PR1 regulates the quiescence of lymphatic vessels by inhibiting laminar shear stress-dependent VEGF-C signaling"

#### Supplementary Figure 1

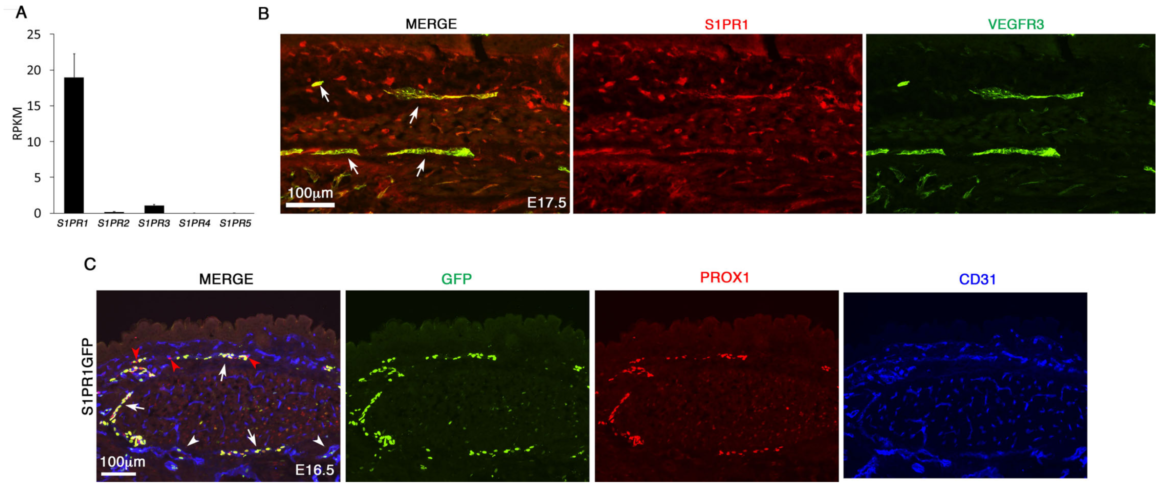


**Supplementary Figure 1: S1PR1 expression and activity are observed in HLECs and mouse LECs.**

*(A) S1PR1 is the most strongly expressed S1P receptor in HLECs.* RNA-seq was performed using primary human LECs and the reads per kilo base of transcript per million mapped reads (RPKM) for the five S1P receptors were plotted. The RPKM values suggest that S1P1 is the most enriched S1P receptor in HLECs.

*(B) S1PR1 is expressed in developing murine lymphatic vessels.* E17.5 embryos were sectioned and analyzed by immunohistochemistry. S1PR1 was expressed in the VEGFR3$^+$ lymphatic vessels of the embryos. Statistics: n=4.

*(C) Heterogeneous S1PR1 activity is observed in murine LECs.* S1PR1-GFP embryos were sectioned and immunostained for GFP. A few GFP$^+$ blood endothelial cells were observed (arrowheads). Most LECs were GFP$^+$ (arrows) although a few GFP$^-$ LECs were also observed (red arrowheads). Statistics: n=4 for both S1PR1GFP and H2B-GFP.

**Supplementary Figure 2**

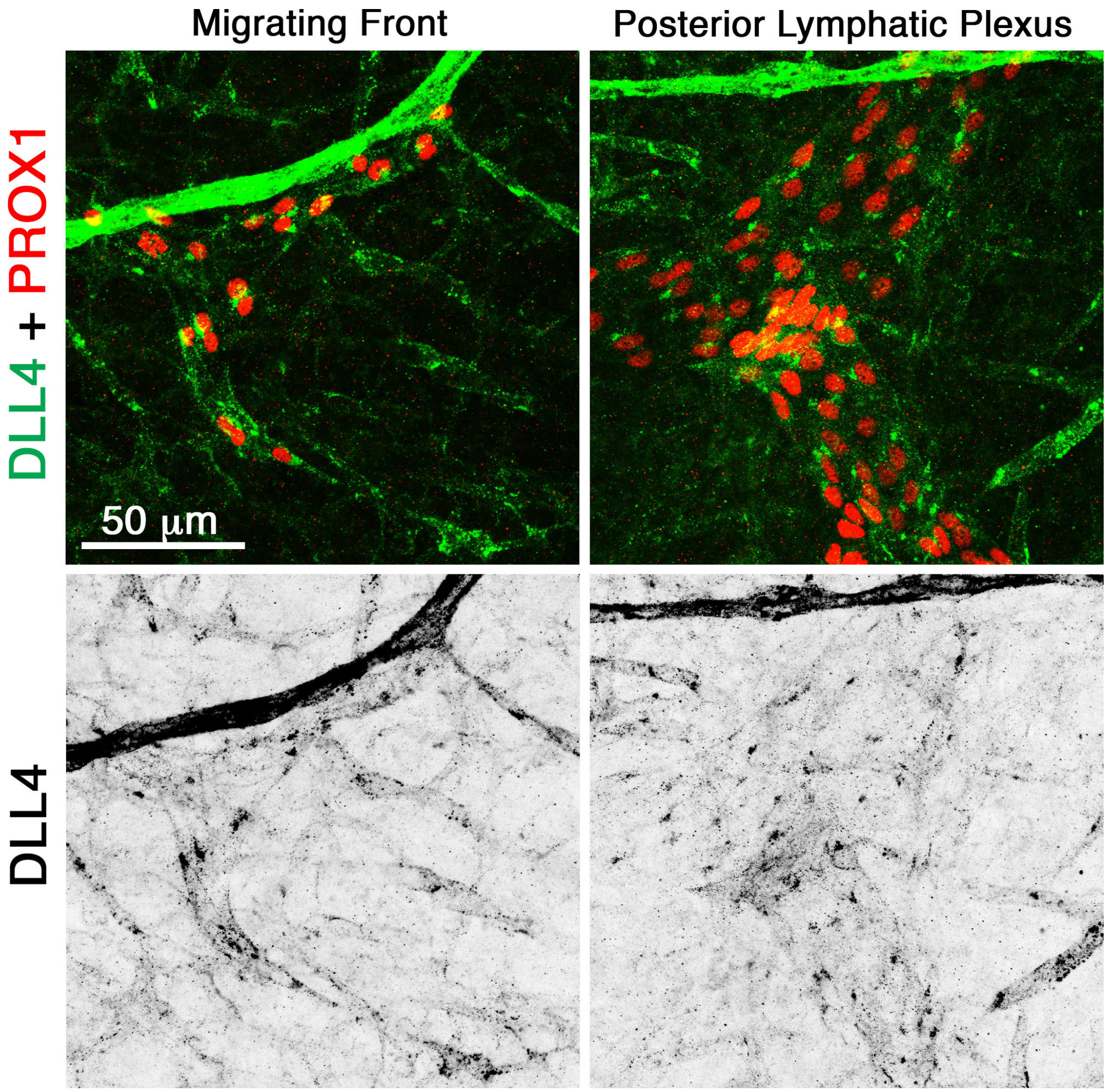


**Supplementary Figure 2: DLL4 expression is enriched in the migrating front of lymphatic vessels.**

Dorsal skins of E16.5 wild type embryos were analyzed for the expression of DLL4. DLL4 was expressed in the migrating front of lymphatic vessels. In contrast, DLL4 expression in the more-mature and quiescent lymphatic vessels was greatly reduced. Statistics: n=5.

**Supplementary Figure 3**

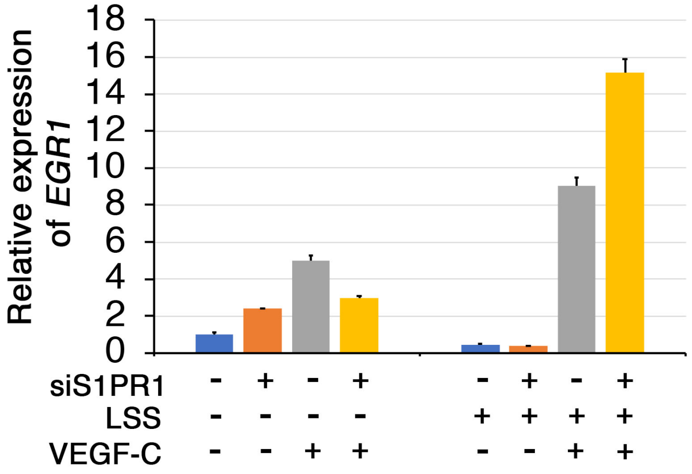


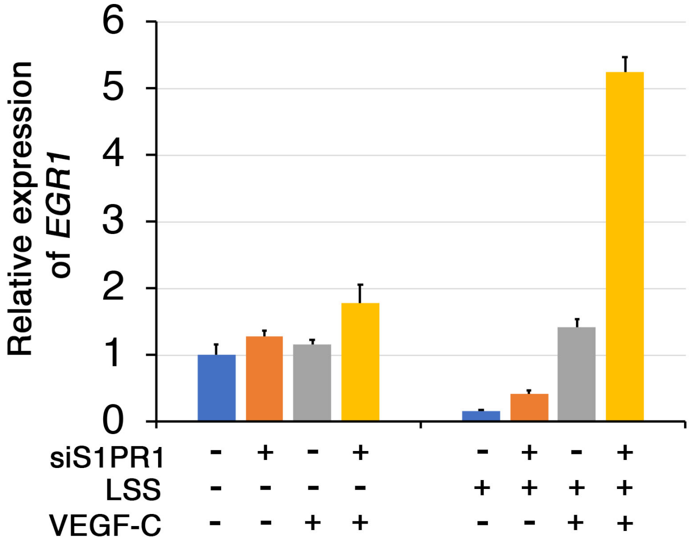


**Supplementary Figure 3: S1PR1 antagonizes LSS/VEGF-C signaling induced expression of EGR1**

VEGF-C induced expression of *EGR1* was dramatically enhanced by LSS and it was further enhanced by siS1PR1.

Statistics: Data from 2 out of 3 independent experiments is presented here. Data from the other experiment is presented in Figure 5C. P values were not presented due to large variability in fold activation of *EGR1* between independent experiments

#### Supplementary Figure 4

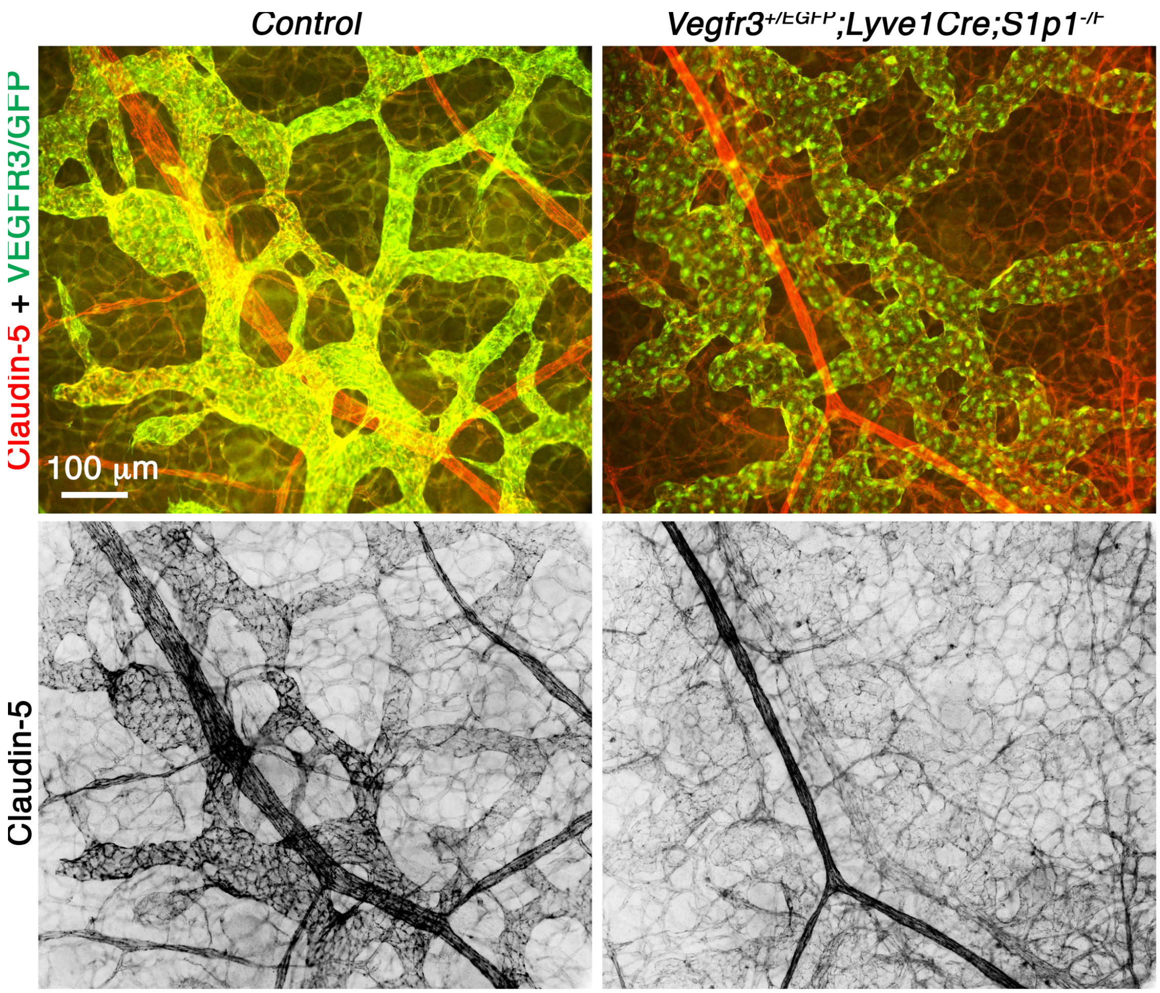


**Supplementary Figure 4: Heterozygosity of *Vegfr3* could not rescue the expression of claudin-5 in embryos lacking S1PR1.**

Claudin-5 was expressed in a gradient manner within the growing lymphatic vessels of control embryos with weaker expression in the migrating tips and stronger expression in the vessels behind. Claudin-5 expression was uniformly downregulated in the lymphatic vessels of *Lyve1-Cre;S1r1*$^{-/f}$ embryos lacking one allele of *Vegfr3*. Statistics: n=5 for control embryos and n=4 for mutant embryos.

### Supplementary Figure 5

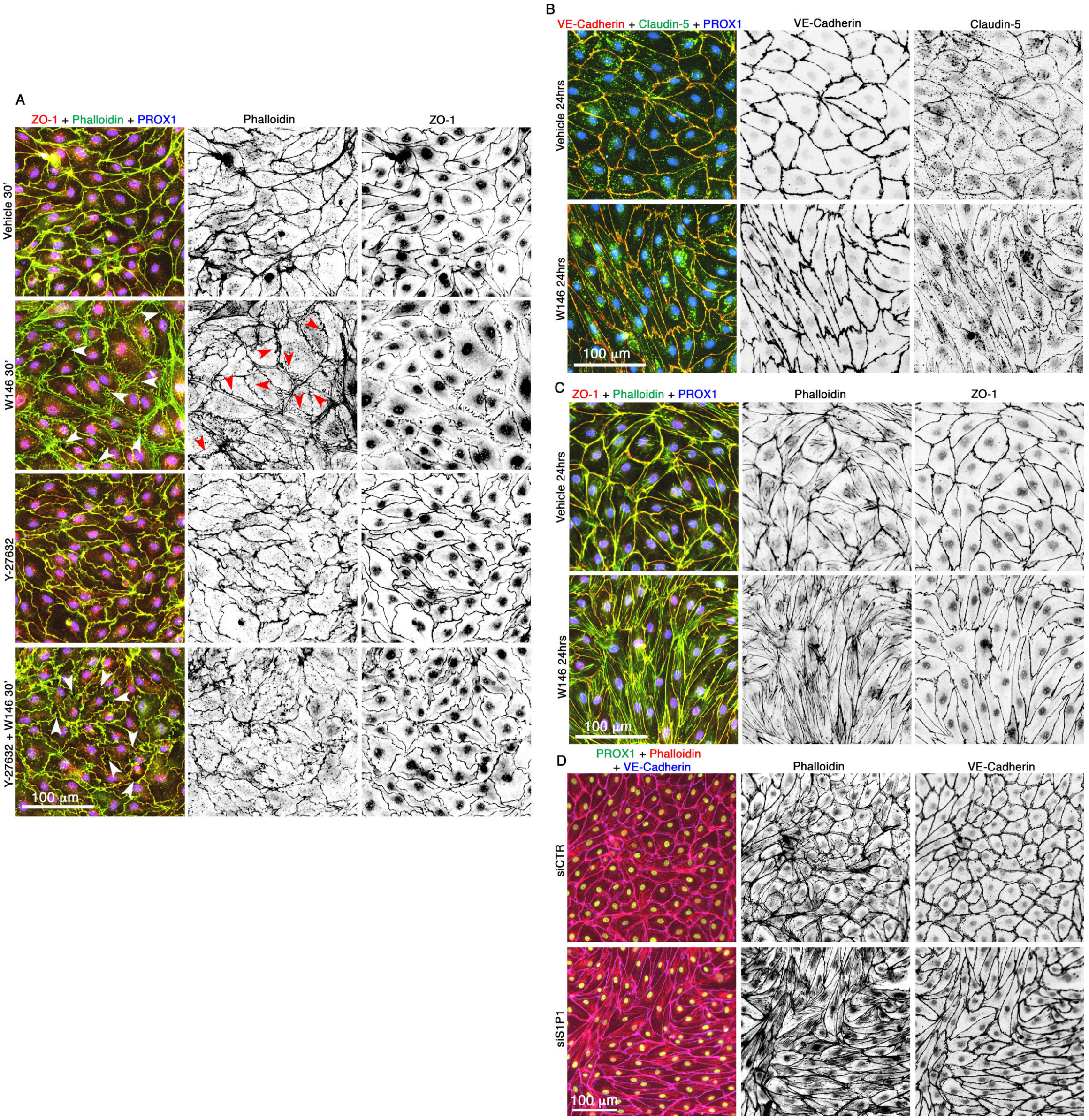

**Supplementary Figure 5: Inhibition of S1PR1 promotes cell junctional and cytoskeletal defects in HLECs.**

*(A) S1PR1 antagonist W146 triggers cytoskeletal reorganization in HLECs, which could be prevented by the inhibition of ROCK.* Phalloidin staining revealed the presence of actin filaments along the periphery of cells (cortical actin) in vehicle treated HLECs. Tight junction molecule ZO-1 was uniformly expressed along the cell membrane. Treatment of HLECs with W146 for 30 minutes resulted in the formation of radial actin bundles, which were aligned perpendicular to cell membrane (red arrowheads). W146 treatment also resulted in the discontinuous expression pattern of ZO-1. Treatment of HLECs with the ROCK inhibitor Y-227632 for 6 hours promoted cell shape change. However, Y-227632 treatment did not cause any obvious defects in the expression of actin or ZO-1. Pretreatment of HLECs with Y-227632 for 6 hours dramatically inhibited the formation of radial filaments upon W146 treatment. White arrowheads point to the intracellular gaps.

*(B-D) Prolonged inhibition of S1PR1 results in the formation of stress fibers and cell elongation.* (B, C) Treatment with W146 for 24 hours resulted in elongated HLECs (B) in which stress fibers could be seen traversing the cytoplasm (C). No obvious defects were observed in the expressions of VE-Cadherin, claudin-5 or ZO-1. (D) Knockdown of S1PR1 from HLECs also resulted in the elongated cells with numerous stress fibers.

Statistics: n=3 for all experiments.

**Supplementary Figure 6**

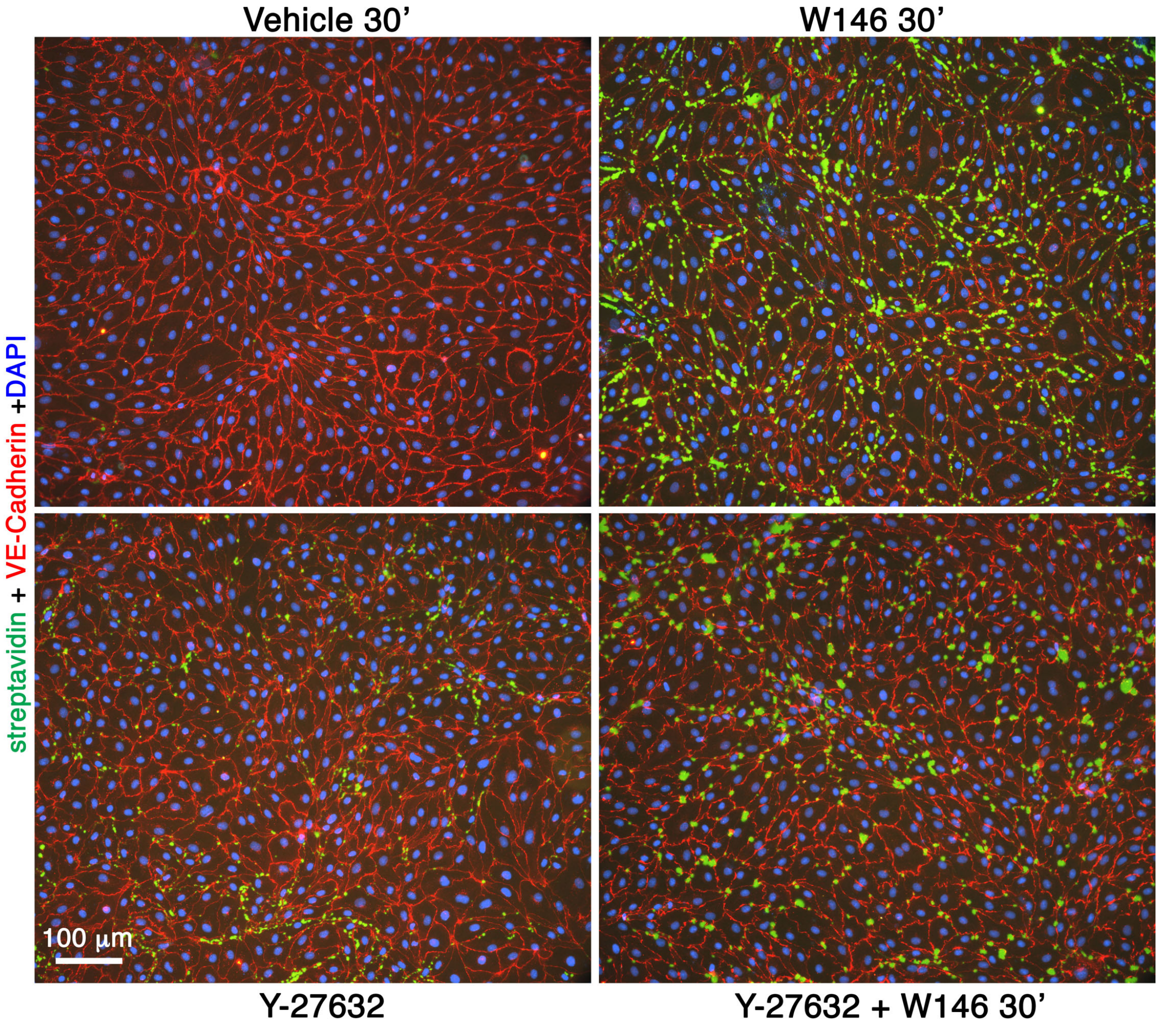


**Supplementary Figure 6: W146 increases the permeability of HLECs and it could not be ameliorated by the inhibition of ROCK.**

Confluent HLECs were treated with W146 for 30 minutes with or without pretreatment with ROCK inhibitor Y-227632 for 6 hours. Intracellular gaps (green signals) were increased by W146. Y-227632 also modestly increased the intracellular gaps. Y-227632 was also not able to prevent the formation of intracellular gaps by W146.
Statistics: n=3.
